# A Simple Method to Distinguish Active and Inactive Aptamers by Analyzing the Ruggedness of the Aptamer Free Energy Landscape

**DOI:** 10.64898/2026.08.26.747184

**Authors:** Gopalakrishnan Subramanian, William Thiel, Rahul Singh

**Author notes:** Corresponding author, Department of Computer Science, The University of Iowa, Iowa City, Iowa, USA.

## Abstract

Aptamers are structured nucleic acid ligands capable of high-affinity, high-specificity molecular recognition generated using variations of the SELEX (Systematic Evolution of Ligands by Exponential Enrichment) process. However, SELEX often produces sequences that enrich yet may lack binding efficacy. We propose a measure called the Ruggedness Composite Index (RCI) along with a method for computing it, that can be used to distinguish binding-competent (“active”) aptamers from weak or non-binding (“inactive”) aptamers. Given a set of aptamers, RCI incorporates information on their fragmentation (landscape partitioning), basin entropy (metastable state distribution), cumulative density irregularity (non-uniform occupancy), and structural–energy correlation length (structure– energy coupling scale). We test whether secondary-structure folding energy landscape topology distinguishes active from inactive aptamers using a multiscale level set framework across six datasets. Active aptamers show lower RCI values and occupy smoother, funnel-like conformational spaces, while inactive aptamers show higher RCI values, reflecting fragmented, high-entropy landscapes. By contrast, classical thermodynamic features, such as minimum free energy, show limited discrimination between active and inactive aptamers. In all datasets, sequences that exhibit enrichment which is not monotonic but lack specificity exhibit elevated ruggedness, indicating landscape topology can predict non-specific enrichment. These results indicate that folding landscape organization can be used as a predictor of aptamer activity and establish RCI as a simple, mechanistically interpretable measure for improving candidate prioritization, especially in therapeutic aptamer discovery.

## INTRODUCTION AND BACKGROUND

Aptamers are short nucleic acid sequences capable of binding molecular targets with high specificity and have attracted interest as therapeutic and diagnostic agents. Identification of aptamers for specific targets most commonly relies on Systematic Evolution of Ligands by Exponential Enrichment (SELEX), an iterative selection process in which pools of sequences are progressively enriched based on target binding.^1^ Despite its conceptual simplicity, SELEX often yields large numbers of candidate sequences whose apparent enrichment does not translate into effective or reproducible activity.^2,3^ This disconnect between enrichment and downstream binding performance remains a central challenge in aptamer research and application, as it necessitates extensive experimental validation to distinguish active aptamers from false positives, thereby increasing experimental cost and prolonging development timelines. Inactive aptamer sequences may initially amplify due to amplification bias,^4,5^ non-specific interactions,^6^ or structural features that confer replication advantages rather than improve affinity.^7^ Consequently, the exclusive use of enrichment-based measures for prioritizing candidates is often ineffective. While experimental strategies such as counterselection and increased selection pressure ^8^ can mitigate such issues, the availability of efficient computational methods that can distinguish between active and inactive aptamers, is limited at the current state-of-the-art.

### Motivating the proposed approach

Increasing evidence suggests that aptamer function is governed not by a single static structure, but by an ensemble of interconverting conformations.^9–15^ In this paper, our focus is on the related (to aptamer function) notion of aptamer biophysical activity. We define an active aptamer as a binding-competent sequence capable of adopting target-specific conformations with high affinity under selection conditions. In contrast, inactive aptamers refer to weak-binding or non-binding sequences.

Enrichment which is not monotonic during selection may not reliably indicate activity consistency; for instance, sequences with unstable or highly partitioned ensembles might amplify due to kinetic or amplification biases, yet fail to sustain consistent target recognition when removed from the selection context.^16–17^ Ensemble properties such as diversity, base-pairing probabilities, conformational switching, and mutational sensitivity have been shown to influence binding and specificity, particularly when active conformations are not the minimum free-energy state or are stabilized upon ligand binding.^18–22^ Several computational approaches have incorporated ensemble-level descriptors, including ensemble entropy,^23^ base-pairing probabilities,^24^ and structural clustering.^25^ However, most existing methods, including machine learning and generative methods,^6,19^ treat these characteristics as “features” and do not mechanistically capture how conformational states are organized and connected across the folding energy landscape.^26–29^

In this context, properties such as basin dominance (the extent to which a single low-free-energy structural basin contains a large fraction of the ensemble population), fragmentation (the degree to which the landscape is partitioned into multiple low-population basins separated by energetic barriers), and *ΔG*-dependent connectivity (the extent to which structural basins are thermodynamically and kinetically connected as a function of free-energy differences) provide a more explicit description of folding landscape organization. Arguably, these features are particularly relevant for distinguishing sequences that can reliably adopt binding-competent conformations from those that cannot; yet they have not been systematically explored in the context of aptamer selection.

We propose characterizing the topological properties of RNA folding energy landscapes and employing this characterization to model aptamer activity. To this end, we introduce a measure called the Ruggedness Composite Index (RCI), where higher RCI values indicate more rugged, fragmented, and heterogeneous folding landscapes that correspond to inactive aptamers, whereas lower RCI values denote smoother, funnel-like landscapes indicative of binding-competent, active aptamers. To our knowledge, this work presents the first multiscale topological characterization of aptamer folding ruggedness, establishing folding landscape organization as a general and mechanistically interpretable determinant of aptamer activity across diverse experimental selection settings.

We analyze multiple high-throughput SELEX datasets spanning distinct targets and selection strategies to evaluate whether folding landscape organization provides information complementary to enrichment-based approaches for prioritizing candidate sequences. By examining enrichment behavior alongside RCI values, we identify sequences that enrich early yet exhibit high RCI, as well as modestly enriched sequences with lower RCI. Sequences with high RCI are disproportionately likely to lose enrichment in later selection rounds, whereas those with lower ruggedness exhibit more stable enrichment trajectories, suggesting that folding landscape organization can help distinguish initial short-lived enrichment from sustained active enrichment.

## EXPERIMENTS AND RESULTS

To assess the RCI framework, we conducted four distinct analyses using six aptamer datasets— ASSET, IL-10, VEGF, Chloramphenicol, Dengue, and Thrombin—which represent a broad range of target types and selection methods. First, we mapped and measured the folding landscapes of active and inactive aptamers through a level-set framework, quantifying these landscapes using measures like fragmentation, basin entropy, and cumulative density. Next, we examined how RCI aligns with early HT-SELEX enrichment stability across the ASSET and IL-10 datasets and benchmarked its performance against traditional thermodynamic descriptors. Finally, we established a straightforward RCI-threshold rule to predict individual aptamer activity, confirming its accuracy through both resubstitution and leave-one-out cross-validation.

### Case Studies on Level-Set Analysis of Folding Landscapes

In the context of RNA folding, the “landscape” is an energy landscape in which each secondary structure corresponds to a point, and and its Gibbs free energy, relative to the unfolded state, defines its height. We analyzed the RNA folding landscape using the concept of level sets.^30–34^ Intuitively, the level set of a function can be thought of as the collection of all points in its domain that share the same value or that lie below (or above) a specified threshold. In the context of an energy landscape, the level sets can be visualized using an energy topographic map with contour lines (Figure 1). Each contour line represents a level set of constant energy, connecting all points in the domain that have the same energy value (for example, *E* = 10 units). The contours trace regions of equal energy across the landscape. A *thresholded energy level set* is analogously defined by considering all points whose energy is less than or equal to a specified (threshold) value. As the energy threshold is changed, progressively larger portions of the energy landscape become included or excluded corresponding to larger or smaller threshold values. In this setting, for our context, a level set consists of all structures whose free energy lies within a specified threshold above the minimum free energy.

**Figure 1.**
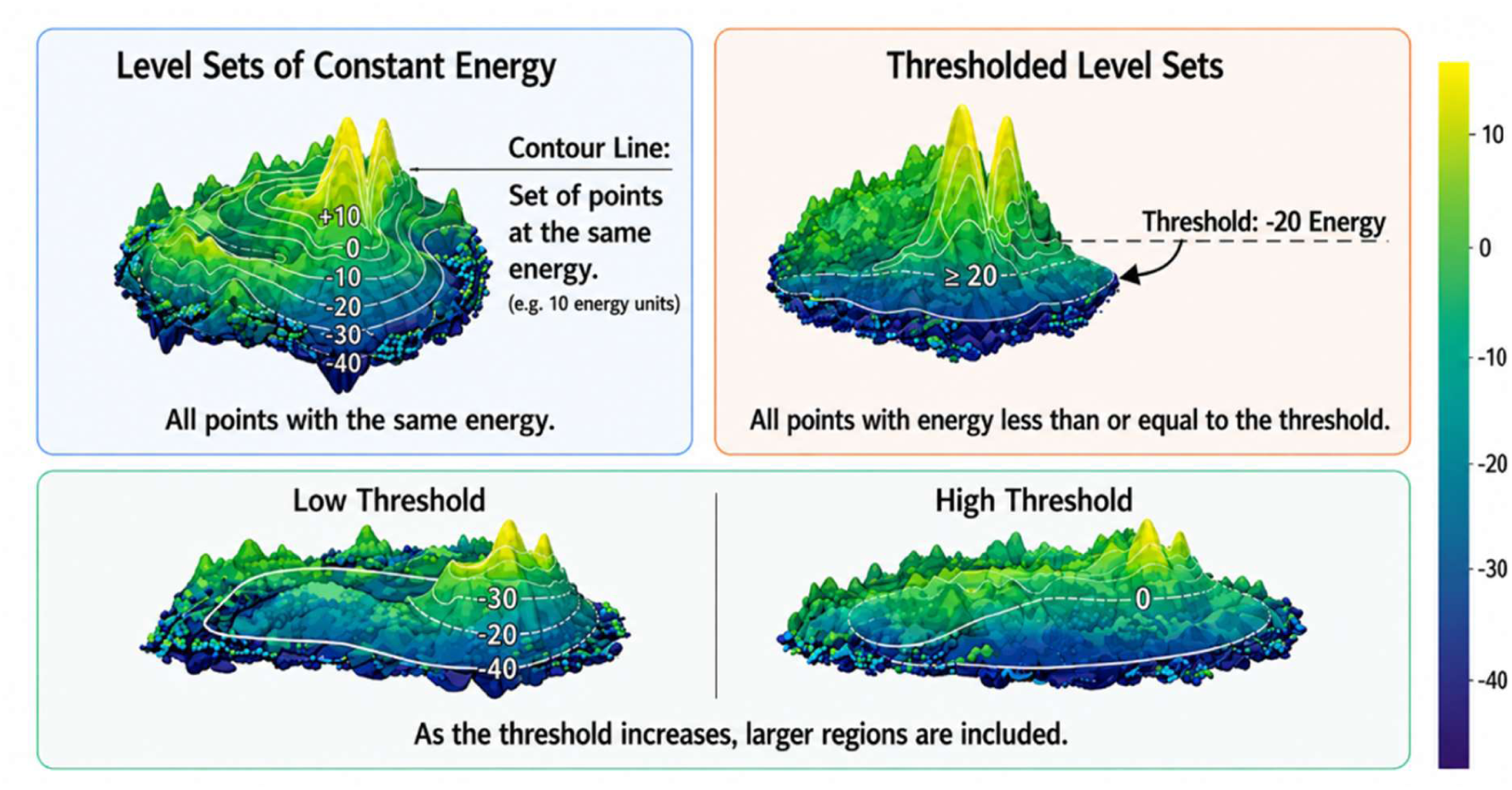
Energy level sets illustrated within a representative folding energy landscape. Level sets of constant energy are shown as contour lines (Top Left), where each contour represents the set of points with the same energy value (e.g., a contour corresponding to increments of +10 energy units). In contrast, thresholded level sets (Top Right) include all points with energy less than or equal to a specified threshold, illustrated here for a threshold of −20 energy units. The lower panels demonstrate how the included region changes with the threshold: at a low threshold (Lower Left), only the deepest portions of the landscape are included, whereas as the threshold increases, progressively larger regions of the landscape become part of the level set. At a high threshold (Lower Right), most of the landscape is encompassed. These examples highlight how level sets can be used to describe the organization of basins and connectivity within the underlying energy landscape.

To visualize folding organization at the ensemble level, three-dimensional free energy landscapes were constructed for all six aptamer datasets (ASSET, IL_10, VEGF, Chloramphenicol, Dengue, and Thrombin).^14,35–39^ Example landscapes are shown in Figures 2A–Figure 2F, with full visualizations provided in Figures S1–S6 and the details describing landscape construction provided in Materials and Methods. These data are described in detail in Dataset Compilation and Activity Definitions section (See Materials and Methods). Each panel shows a 3D map of how an aptamer folded, where the height represents energy and the surface shape reflects how many alternative structures were available. Each panel shows a 3D map of how an aptamer folded, where the height represents energy and the surface shape reflects how many alternative structures were available. Binding-competent aptamers were classified as being “active” and weak or non-binding aptamers were classified as “inactive”. Active aptamers tended to have smoother, funnel-like landscapes that guided folding into a stable structure, whereas inactive aptamers displayed rough, jagged landscapes with many competing states, indicating less stable and less specific folding behavior.

**Figure 2.**
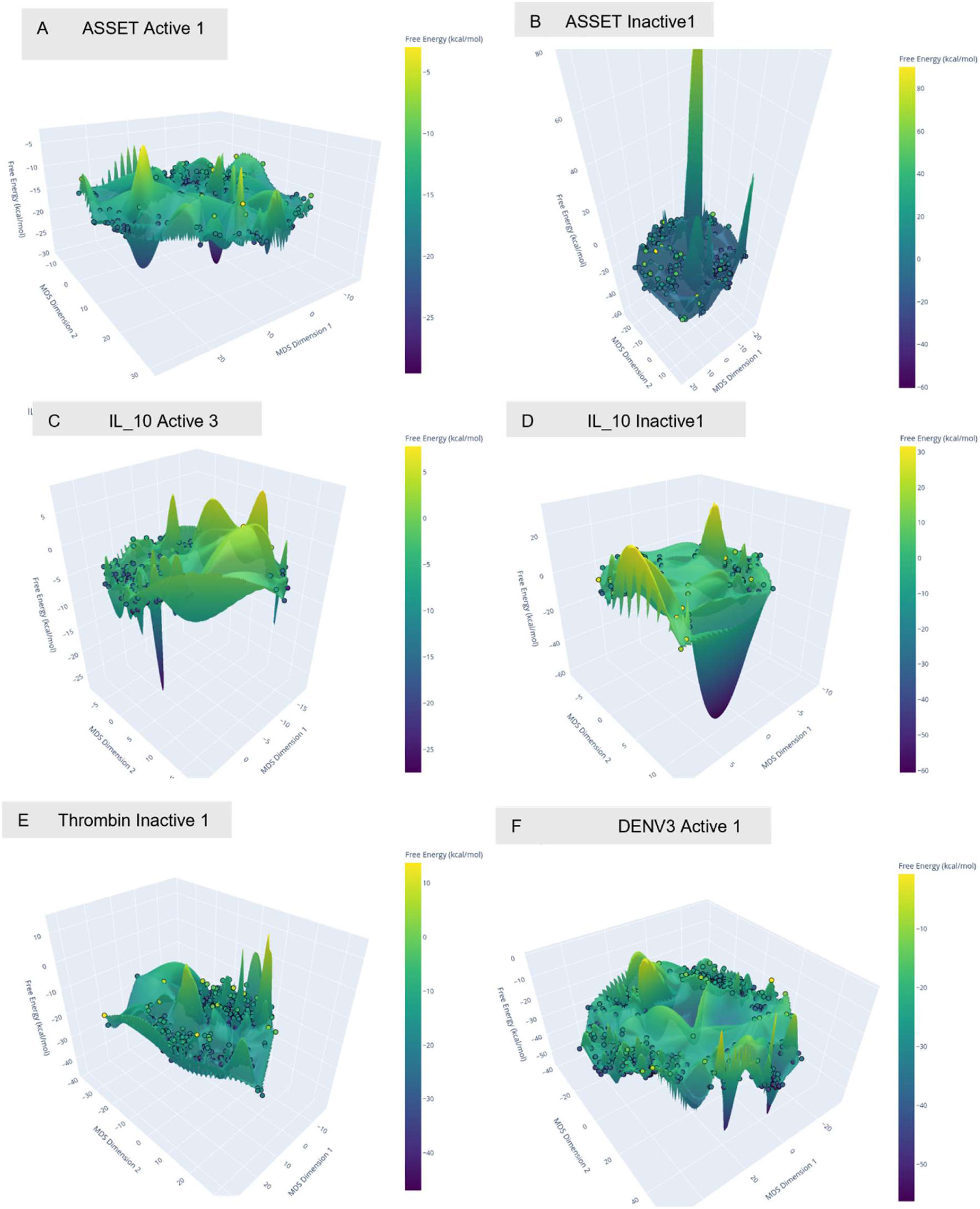
Representative Aptamer Free Energy Landscapes of Active and Inactive Aptamers. (A) ASSET Active_1 Aptamer, (B) ASSET Inactive_1 Aptamer, (C) IL_10 Active_3 Aptamer, (D) IL_10 Inactive1 Aptamer, (E) Thrombin2 Inactive1 Aptamer, (F) Dengue DENV-3_Active1 Aptamer

We applied the level set framework^30–34^ to track and analyze how conformational connectivity evolved as progressively higher-energy states became accessible and used the resulting changes in connectivity to characterize aspects of landscape ruggedness versus smoothness. Smooth (low ruggedness) landscapes maintained coherent, well-connected basins as energy increased, whereas rugged landscapes fragmented into multiple disconnected regions with heterogeneous structural distributions. Building on this energy-stratified landscape representation, structures for each sequence were grouped according to *ΔG* above the minimum free energy and connectivity graphs were constructed at each energy threshold to capture how conformational connectivity emerged across the landscape. Three *ΔG*-dependent measures were computed: (i) number of connected components (fragmentation), (ii) basin entropy (distribution of structural mass across basins), and (iii) cumulative structural density (Panels i, ii, iii in Figures 3-5). Together, these measures quantified the degree of fragmentation and heterogeneity across energy levels, providing the basis for assessing folding landscape ruggedness.

**Figure 3.**
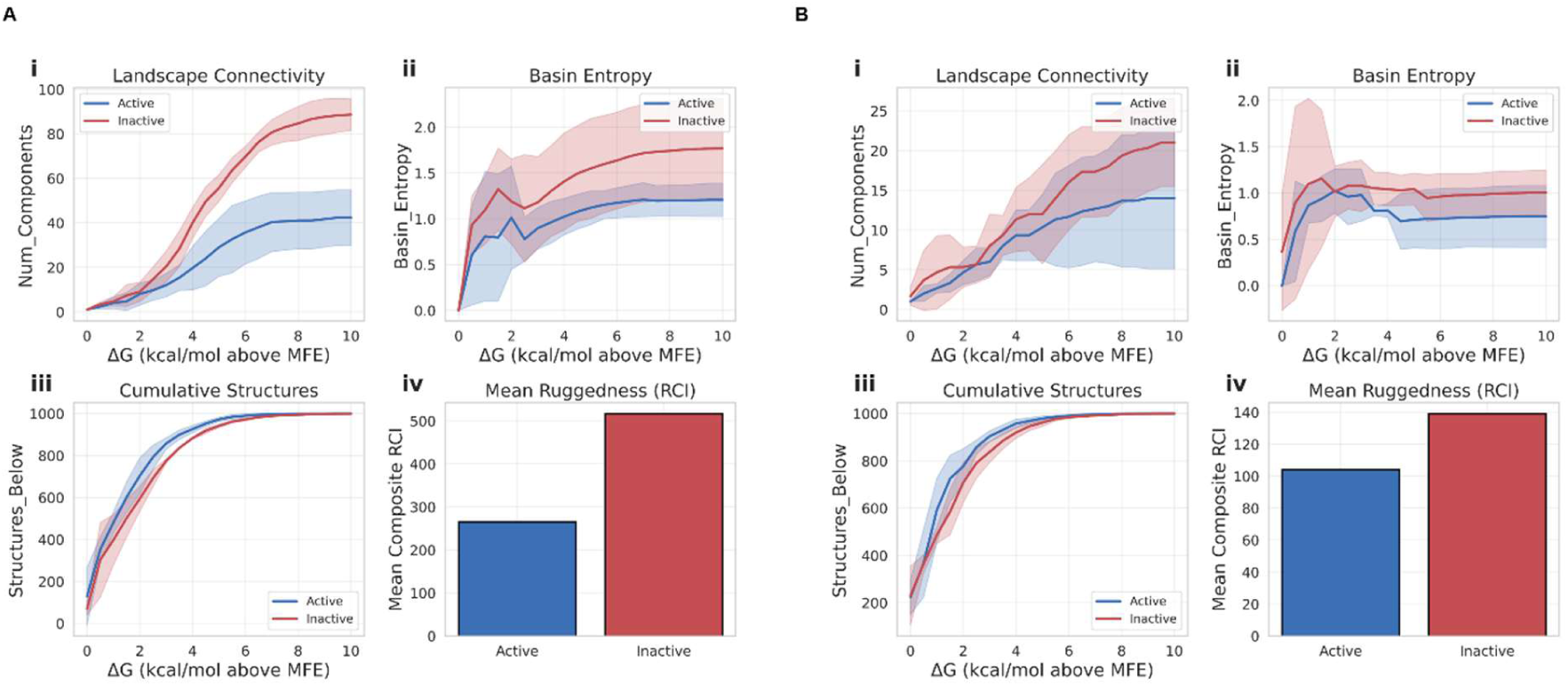
Comparison of structural energy landscape ensemble properties between active and inactive groups of aptamers from the ASSET and IL_10 data sets. (A) **ASSET Data Set:** Landscape connectivity (Panel i) increased more strongly in inactive aptamers (∼81 vs ∼31 components at *ΔG* = 10 kcal/mol; ∼2.6-fold). Basin entropy (Panel ii) was higher in inactive sequences (∼1.93 vs ∼1.10; ∼75% increase), indicating broader conformational diversity. Active aptamers accumulated low-energy structures (Panel iii) more rapidly at early thresholds (0–3 kcal/mol), consistent with enrichment of active folds. Mean ruggedness (RCI) (Panel iv) was higher in inactive sequences (∼488 vs ∼260; ∼1.9-fold), reflecting more complex landscapes. Shaded regions indicate variability (mean ± SD or CI). (B) **IL_10 Data Set:** Landscape connectivity (Panel i) was higher in inactive aptamers (∼22 vs ∼12 at *ΔG* = 10 kcal/mol; ∼1.8-fold). Basin entropy (Panel ii) was modestly elevated in inactive sequences, converging near *ΔG* = 10 (∼0.91 vs ∼0.88). Active aptamers accumulated low-energy structures earlier (Panel iii), while ruggedness (RCI) (Panel iv) remained higher in inactive sequences (∼138 vs ∼100; ∼1.4-fold). Shaded regions indicate variability (mean ± SD or CI). Overall, inactive IL_10 aptamers retained greater landscape complexity, whereas active sequences exhibited smoother, more energetically focused ensembles consistent with SELEX enrichment

**Figure 4.**
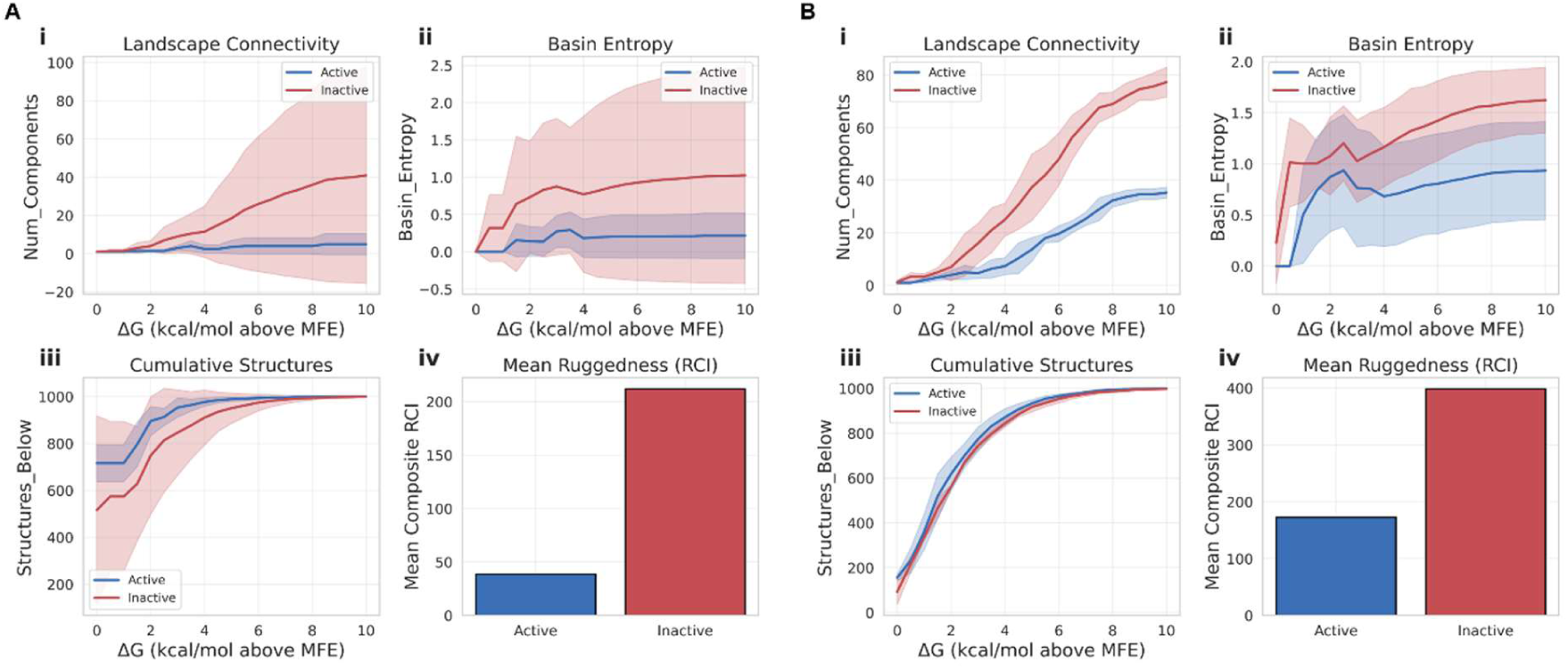
Comparison of structural energy landscape ensemble properties between active and inactive groups of aptamers from VEGF and Chloramphenicol data sets. (A) **VEGF Data Set:** Landscape connectivity (Panel i) was higher in inactive aptamers (∼34 vs ∼8 components at *ΔG* = 10 kcal/mol; ∼4.3-fold), indicating strong structural fragmentation. Basin entropy (Panel ii) was elevated in inactive sequences (∼1.03 vs ∼0.25; ∼4.1-fold), reflecting broader conformational diversity. Active aptamers accumulated low-energy structures (Panel iii) more rapidly, reaching ∼990 by *ΔG* ≈ 4, while inactive sequences converged later. Mean ruggedness (RCI) (Panel iv) was higher in inactive sequences (∼210 vs ∼40; ∼5.3-fold), indicating highly complex landscapes. Shaded regions represent variability (mean ± SD or CI). Overall, VEGF shows the strongest separation between active and inactive pools, consistent with effective SELEX enrichment. (B) **Chloramphenicol Data Set:** Landscape connectivity (Panel i) was higher in inactive aptamers (∼70 vs ∼36 at *ΔG* = 10 kcal/mol; ∼1.9-fold). Basin entropy (Panel ii) remained elevated in inactive sequences (∼1.55 vs ∼1.05; ∼1.5-fold). Cumulative structures (Panel iii) increased for both groups, with slightly faster early accumulation in active aptamers and convergence near ∼1000 by *ΔG* ≈ 7–8. Mean ruggedness (RCI) (Panel iv) was higher in inactive sequences (∼360 vs ∼180; ∼2.0-fold). Shaded regions represent variability (mean ± SD or CI).

**Figure 5.**
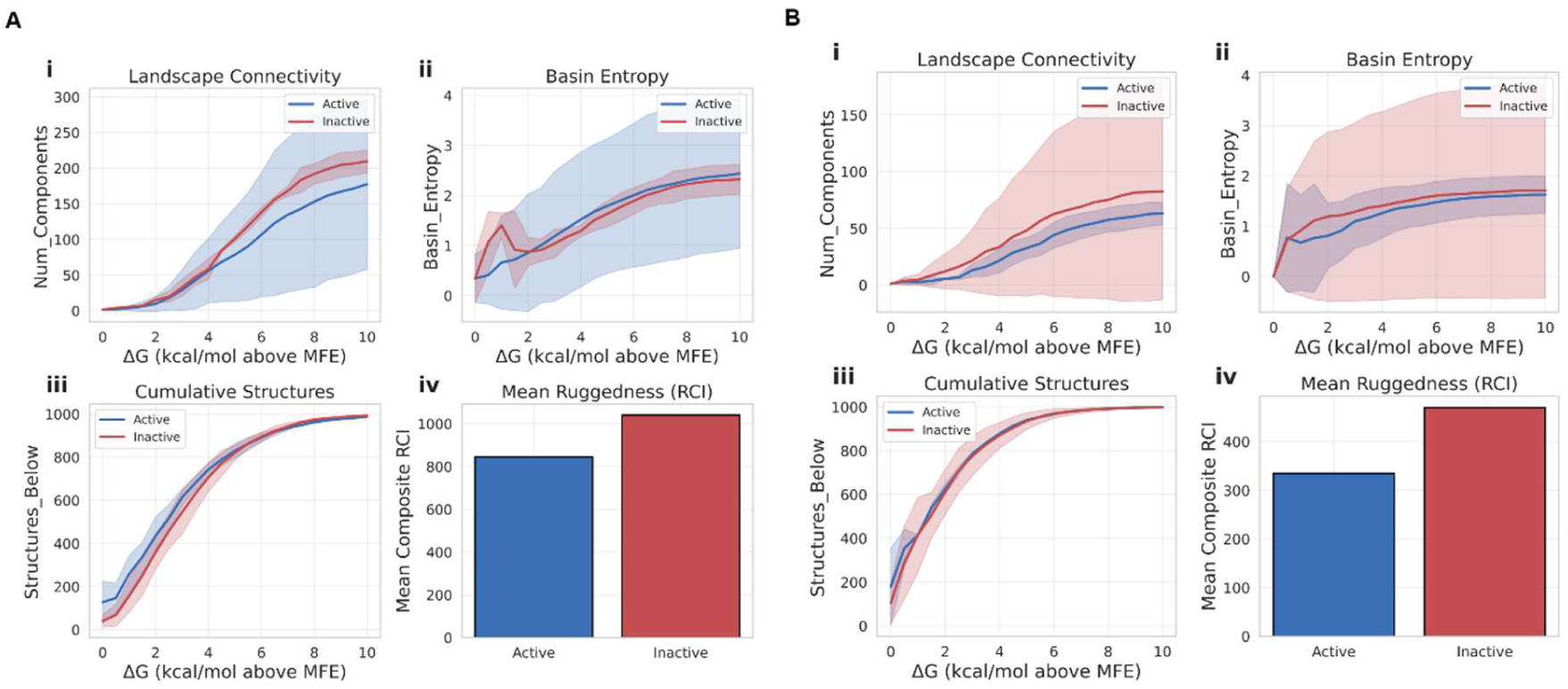
Comparison of structural energy landscape ensemble properties between active and inactive groups of aptamers from Dengue and Thrombin data sets. (A) Dengue Data Set: Landscape connectivity (Panel i) was higher in inactive aptamers (∼225 vs ∼170 at *ΔG* = 10 kcal/mol; ∼1.3-fold), indicating moderate structural fragmentation. Basin entropy (Panel ii) was slightly elevated in inactive sequences (∼2.5 vs ∼2.4; ∼1.05-fold), suggesting similar but marginally broader conformational diversity. Cumulative structures (Panel iii) accumulated rapidly in both groups, with a small early lead in active aptamers (∼550 vs ∼500 at *ΔG* ≈ 2) and convergence near ∼1000 by *ΔG* ≈ 8– 10. Mean ruggedness (RCI) (Panel iv) was higher in inactive sequences (∼1100 vs ∼830; ∼1.3-fold), indicating more complex landscapes. Shaded regions represent variability (mean ± SD or CI). **(B) Thrombin Data Set:** Landscape connectivity (Panel i) was higher in inactive aptamers (∼90 vs ∼62 at *ΔG* = 10 kcal/mol; ∼1.5-fold). Basin entropy (Panel ii) was modestly elevated in inactive sequences (∼1.7 vs ∼1.6; ∼1.1-fold). Cumulative structures (Panel iii) increased rapidly for both groups, with a slight early advantage in active aptamers (∼800 vs ∼750 at *ΔG* ≈ 3) and convergence near ∼1000 by *ΔG* ≈ 7–8. Mean ruggedness (RCI) (Panel iv) was higher in inactive sequences (∼480 vs ∼330; ∼1.45-fold). Shaded regions represent variability (mean ± SD or CI).

Active aptamers consistently exhibited smoother energy landscapes, defined quantitatively by lower fragmentation (fewer connected components at a given *ΔG*), lower basin entropy (more concentrated structural occupancy), and more gradual changes in these measures as *ΔG* increased. As higher-energy states became accessible, active sequences showed a slow, continuous growth in connected components and moderate increases in basin entropy, consistent with cooperative, funnel-like organization. In contrast, inactive sequences fragmented rapidly at low *ΔG* values and displayed higher basin entropy across energy thresholds, indicating dispersal across competing conformations. These trends were summarized using the RCI (Figures 3-5).

Across all six datasets, at the final round of SELEX, inactive aptamers exhibited higher mean RCI values than active aptamers. For each dataset the average group-wise RCI differences between the average active vs inactive aptamers exceeded within-group variability, yielding minimal overlap between distributions despite differences in sequence length, target type, and experimental protocol (Figure 3-5, Panel iv)

### Early Enrichment Behavior and RCI values

We used sequences from all six data sets while comparing RCI values between active aptamers to inactive aptamers. The ASSET and IL_10 data sets provided full sets of aptamers with associated round to round SELEX enrichment data, which the other four data sets did not include. Consequently, we used the ASSET and IL_10 data for exploring the connection between RCI and early enrichment behavior. In particular, to evaluate how folding landscape organization related to SELEX enrichment dynamics, we examined round-by-round enrichment data from the ASSET HT-SELEX and IL_10 datasets and computed the RCI values of aptamers which demonstrate initial short-lived enrichment against active aptamers from each data set. Almost all sequences which displayed significant early enrichment (*e.g.,* Round 4 relative to Round 3 for the ASSET aptamers and Round 4 relative to Round 2 for the IL_10 aptamers) failed to retain enrichment in later rounds (*e.g.,* Round 9 relative to Round 8 for the ASSET aptamers and Round 5 relative to Round 4 for the IL_10 aptamers) and were ultimately classified as inactive. Despite initial short-lived enrichment signatures, these sequences consistently exhibited elevated RCI values comparable to inactive aptamers and significantly higher than final-round active sequences. (Figure 6 and Figure S7).

**Figure 6.**
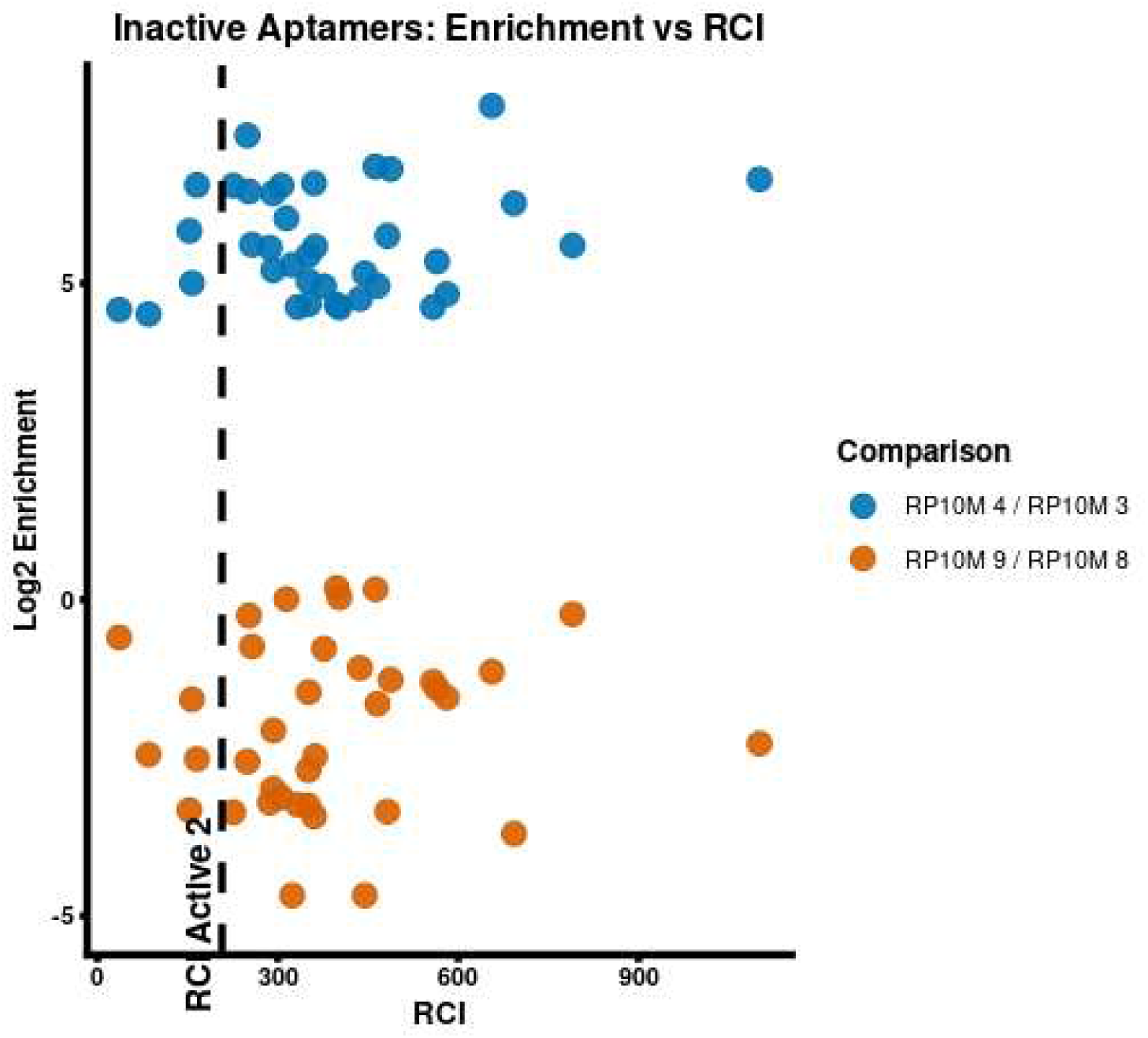
Round-to-round enrichment behavior and folding landscape Ruggedness Composite Index (RCI) of ASSET Inactive aptamers. The figure shows sequence-level enrichment ratios across early and late HT-SELEX rounds (Blue Points represent Round 4/ Round 3 and Orange Points represent Round 9/ Round 8) plotted against the RCI computed from sequence-derived folding energy landscapes with low RCI value of ASSET Active2 aptamer shown as reference. A substantial proportion of sequences exhibit significant positive enrichment at early rounds yet fail to retain enrichment at later rounds and are ultimately classified as inactive based on final ASSET specificity criteria. These early-enriched but inactive sequences consistently display elevated RCI values, indicative of fragmented and high-entropy folding landscapes, whereas sequences that remain enriched through later rounds and are classified as active exhibit lower RCI values. The figure illustrates that landscape ruggedness provides an independent discriminator of activity outcome beyond enrichment trajectories alone

### Analysis of the ability of Thermodynamic and Structural Features to Distinguish Active from Inactive Aptamers

To evaluate whether thermodynamic stability measures and higher-order structural descriptors could distinguish active aptamers from inactive ones, we analyzed experimentally derived RNA aptamer datasets representing different targets and SELEX selection protocols (Table S1). Two datasets, namely the ASSET HT-SELEX aptamer dataset^30^ and the IL_10 aptamer dataset,^14^ contained experimentally determined activity labels that classified sequences as active or inactive.

Active ASSET aptamers (n = 115) exhibited a modest shift toward more negative Minimum Free Energy (MFE) values compared to inactive aptamers, with substantial overlap between the two distributions (Figure S8A, Figure S8B). The median MFE of active sequences was approximately −17.0 kcal·mol⁻¹ (IQR ≈ −18.5 to −15.8), whereas inactive sequences showed a slightly less negative median around −16.0 kcal·mol⁻¹ (IQR ≈ −17.8 to −14.5). Both density and boxplot visualizations indicated that while active aptamers tended to have somewhat lower (more negative) MFEs— suggesting a tendency toward more thermodynamically stable predicted secondary structures—the difference was relatively small and the distributions largely overlapped.

Similarly, active IL_10 aptamers exhibited MFE values that broadly overlapped with inactive sequences (Figure S9 A, Figure S9 B), with no statistically significant difference between the two distributions (e.g., Wilcoxon rank-sum test, *p* > 0.05). By contrast, inactive sequences had a median MFE of approximately –4.0 kcal·mol⁻¹ (IQR ≈ –8.6 to –3.6), indicating comparable central tendencies but somewhat greater dispersion among inactive aptamers. However, substantial distributional overlap remained, indicating that MFE alone could not reliably distinguish active classes based on the MFE analysis for both the ASSET and IL_10 aptamers.

Other descriptors showed limited discriminatory power in distinguishing active and inactive aptamers. Energy gap, ensemble diversity, and number of basins did not differ significantly between the two groups (*p* > 0.5; (Figure S8C–Figure S8H) for ASSET aptamers and Figures S9C-S9H for IL_10 aptamers). Although these features characterized aspects of ensemble flexibility and structural dispersion, they failed to provide clear separation when considered individually. Sequence length (∼55–57 nt) (Figure S8I) and GC content (0.60–0.70) (Figure S8J) were comparable between active and inactive groups for ASSET aptamers and Figure S9I, Figure S9J for IL_10 aptamers), indicating that basic compositional properties did not account for observed activity differences.

Basin size distributions (Figure S8K) and energy gap spectra (Figure S8L) for ASSET aptamers and Figure S9K, Figure S9L for IL_10 aptamers) further revealed that aptamer folding landscapes generally contained multiple metastable conformations separated by modest energetic barriers (up to ∼1.2 kcal·mol⁻¹). Such heterogeneity is typical of RNA systems^40^ and suggests that activity discrimination likely depends on higher-order landscape organization rather than on isolated scalar features.

To visualize structural clustering of active and inactive aptamers, we performed linear (classical MDS) and non-linear (t-SNE, UMAP) dimensionality reduction on this feature space. Across both the ASSET and IL_10 datasets, these lower-dimensional projections showed substantial overlap, yielding only partial separation between active and inactive sequences (Figures S10A–C and S11A–C). These findings highlight the fundamental limitations of traditional scalar features and reinforce the necessity of RCI, which captures multiscale, topological landscape organization rather than isolated measures.

### RCI based Activity Classification of Aptamers

To classify aptamers as Active or Inactive solely from RCI, we established a label-agnostic threshold for each dataset by sorting all RCI values, finding the sparsest interval between the RCI scores distributions, and placing the cutoff at its midpoint. This “sparsest-interval” strategy bypasses the flaws of label-dependent boundaries: mislabeled data cannot skew it and overlapping active/inactive distributions do not disrupt it because it evaluates the complete dataset rather than group edges. Table S2 details individual RCI measures, Ground Truth, and predictions for all 30 sequences, highlighting one or two misclassified aptamers across four datasets (VEGF, Dengue, Thrombin, IL-10).

Resubstitution testing across all six datasets (Table 1) yielded 80.0% pooled accuracy (24/30 correct; MCC = 0.61). Under leave-one-out cross-validation (Table 2), pooled accuracy rose to 83.3% (25/30; MCC = 0.67), maintaining identical results in five datasets (VEGF, Chloramphenicol, Dengue, Thrombin, ASSET) while boosting IL-10 (66.7% → 83.3%). Maintaining—and even improving— accuracy during cross-validation confirms that this threshold avoids overfitting, unlike rules based on group means or extremes.

**Table 1.** RCI-Based Activity Prediction Performance across Datasets (Resubstitution), Sparsest-Interval Threshold. This table summarizes confusion-matrix performance for the RCI predictions in Table S2 against experimental Ground Truth, treating Active as the positive class. The evaluation applies the sparsest-interval threshold—the midpoint of the widest gap in each dataset’s sorted RCI distribution, derived without Ground Truth labels—and uses resubstitution to assess accuracy on the training data. For each dataset, it reports true positives (TP), false positives (FP), true negatives (TN), false negatives (FN), sensitivity [TP/(TP + FN)], specificity [TN/(TN + FP)], accuracy [(TP + TN)/*n*], precision [TP/(TP + FP)], F1 score, and Matthews correlation coefficient (MCC). The final row aggregates raw counts across all six datasets to calculate overall performance measures.

| Data Set | n | TP | FP | TN | FN | Sens. | Spec. | Acc. | Prec. | F1 | MCC |
| --- | --- | --- | --- | --- | --- | --- | --- | --- | --- | --- | --- |
| VEGF | 4 | 2 | 1 | 1 | 0 | 100.0% | 50.0% | 75.0% | 66.7% | 0.80 | 0.58 |
| Chloramphenicol | 6 | 3 | 0 | 3 | 0 | 100.0% | 100.0% | 100.0% | 100.0% | 1.00 | 1.00 |
| Dengue | 4 | 1 | 0 | 2 | 1 | 50.0% | 100.0% | 75.0% | 100.0% | 0.67 | 0.58 |
| Thrombin | 4 | 2 | 1 | 1 | 0 | 100.0% | 50.0% | 75.0% | 66.7% | 0.80 | 0.58 |
| ASSET | 6 | 2 | 0 | 3 | 1 | 66.7% | 100.0% | 83.3% | 100.0% | 0.80 | 0.71 |
| IL-10 | 6 | 3 | 2 | 1 | 0 | 100.0% | 33.3% | 66.7% | 60.0% | 0.75 | 0.45 |
| **All datasets (pooled)** | **30** | **13** | **4** | **11** | **2** | **86.7%** | **73.3%** | **80.0%** | **76.5%** | **0.81** | **0.61** |

**Table 2.** RCI-Based Activity Prediction Performance across Datasets: Resubstitution versus Leave-One-Out Cross-Validation under a Sparsest-Interval Threshold. This table compares classification measures for the sparsest-interval threshold rule (defined in Table S2) across two validation methods. Resubstitution columns display performance when fitting and testing the threshold on all *n* points in a dataset, mirroring Table S2. Leave-One-Out Cross-Validation (LOOCV) columns present performance when the algorithm sequentially holds out each sequence, recalculates the threshold using only the remaining *n* − 1 points, and blindly predicts the held-out aptamer’s class. The measures—sensitivity, specificity, accuracy, precision, F1 score, and MCC—match those in Table 1, with the final row pooling raw counts across all six datasets to recalculate total performance. LOOCV yields identical results to resubstitution for VEGF, Chloramphenicol, Dengue, Thrombin, and ASSET. Conversely, IL-10 accuracy increases under LOOCV (66.7% → 83.3%), demonstrating that resubstitution underestimated its out-of-sample performance. Across all six pooled datasets, LOOCV boosts overall accuracy from 80.0% to 83.3% and raises MCC from 0.61 to 0.67, confirming that the sparsest-interval threshold generalizes effectively under rigorous cross-validation.

| Data Set | n | *Resub.*<br>Sens. | *Resub.*<br>Spec. | *Resub.*<br>Acc. | *Resub.*<br>Prec. | *Resub.*<br>F1 | *Resub.*<br>MC<br>C | *LOOC*<br>V<br>Sens. | *LOOC*<br>V<br>Spec. | *LOOC*<br>V<br>Acc. | *LOOC*<br>V<br>Prec. | *LOOC*<br>V<br>F1 | *LOOC*<br>V<br>MCC |
| --- | --- | --- | --- | --- | --- | --- | --- | --- | --- | --- | --- | --- | --- |
| VEGF | 4 | 100.0<br>% | 50.0% | 75.0% | 66.7% | 0.80 | 0.58 | 100.0<br>% | 50.0% | 75.0% | 66.7% | 0.80 | 0.58 |
| Chloramphenicol | 6 | 100.0<br>% | 100.0<br>% | 100.0<br>% | 100.0<br>% | 1.00 | 1.00 | 100.0<br>% | 100.0<br>% | 100.0<br>% | 100.0<br>% | 1.00 | 1.00 |
| Dengue | 4 | 50.0% | 100.0<br>% | 75.0% | 100.0<br>% | 0.67 | 0.58 | 50.0% | 100.0<br>% | 75.0% | 100.0<br>% | 0.67 | 0.58 |
| Thrombin | 4 | 100.0<br>% | 50.0% | 75.0% | 66.7% | 0.80 | 0.58 | 100.0<br>% | 50.0% | 75.0% | 66.7% | 0.80 | 0.58 |
| ASSET | 6 | 66.7% | 100.0<br>% | 83.3% | 100.0<br>% | 0.80 | 0.71 | 66.7% | 100.0<br>% | 83.3% | 100.0<br>% | 0.80 | 0.71 |
| IL-10 | 6 | 100.0<br>% | 33.3% | 66.7% | 60.0% | 0.75 | 0.45 | 100.0<br>% | 66.7% | 83.3% | 75.0% | 0.86 | 0.71 |
| **All datasets**<br>**(pooled)** | **30** | **86.7%** | **73.3%** | **80.0%** | **76.5%** | **0.81** | **0.61** | **86.7%** | **80.0%** | **83.3%** | **81.2%** | **0.84** | **0.67** |

## DISCUSSION

The methodological framework developed in this study is grounded in a simple premise motivated by biophysics: aptamer function depends not only on the stability of a single predicted structure, but on how the entire folding ensemble is organized across energy scales. Thus, during SELEX, sequences that ultimately prove active must reliably populate binding-competent conformations under equilibrium conditions and maintain structural coherence under environmental and mutational perturbations. From an energy landscape perspective, this implies that active aptamers occupy relatively smooth, funnel-like folding landscapes, defined by a progressive decrement in free energy toward a dominant minimum and a high degree of connectivity among low-energy states, such that conformational probability mass is concentrated within well-connected basins. In contrast, sequences that fold into highly fragmented, high-entropy ensembles distribute structural probability across many competing conformations, reducing the likelihood that any single structure is sufficiently populated to support consistent binding.

Consequently, (quantitatively) characterizing multiscale landscape organization, rather than relying solely on minimum free energy or enrichment values, can help in distinguishing between active and inactive sequences. In essence, our method translates a biophysical intuition—that active folding requires organized conformational accessibility—into a measurable topological descriptor of the folding ensemble.

Classical thermodynamic descriptors such as MFE describe the stability of a single predicted structure but provide limited insight into the broader conformational ensemble. Our results indicate that aptamer activity is more closely associated with folding landscape organization than with individual structural stability. Through our studies, we observed that active aptamers exhibited gradual connectivity among structurally similar conformations as energy thresholds increased, consistent with coherent folding funnels that guide the ensemble toward related states. In contrast, inactive sequences showed rapid fragmentation at low energy thresholds, indicating heterogeneous ensembles in which structurally dissimilar conformations remain energetically competitive and no dominant structural basin emerges. The level set framework revealed these differences across energy scales. Active aptamers displayed lower basin entropy and longer structural–energy correlation lengths, suggesting that sequence constrains channel folding toward organized regions of conformational space where active motifs remain structurally accessible. The persistence of these patterns across datasets with differing sequence lengths, targets, and experimental conditions underlines the folding-landscape topology to be a general descriptor of aptamer activity and not a dataset-specific artifact.

This landscape-centric perspective can be employed to complement existing predictive approaches. Traditional MFE- or motif-based analyses neglect ensemble organization, while machine learning methods^6,19^ often require target-specific training and may lack mechanistic interpretability. In contrast, RCI is directly derived from thermodynamic ensembles and provides a physically interpretable description of folding topology that generalizes across systems without retraining. It is not a replacement for enrichment analysis or experimental validation, but an independent structural descriptor.

Importantly, integration with SELEX dynamics demonstrated practical relevance. In the ASSET HT-SELEX dataset, sequences that enriched early yet failed to retain specificity exhibited elevated RCI values compared to active aptamers, whereas sequences with sustained enrichment showed lower ruggedness, as indicated by lower RCI values. These observations suggest that landscape topology may help identify enrichment-driven, non-specific aptamers prior to final selection convergence and support early-stage candidate triage. Aptamer sequences with rugged, fragmented folding landscapes may initially amplify due to non-specific interactions or amplification bias but lack the structural organization necessary for sustained target-competent folding. Conversely, smoother landscapes are associated with reproducible active enrichment. Thus, RCI provides complementary information to enrichment magnitude and offers a sequence-derived indicator capable of identifying enrichment-driven non-specific aptamers prior to final selection convergence.

The sparsest-interval method of identifying the RCI threshold for activity classification of aptamers requires no prior assumptions about class boundaries. This feature provides a key methodological edge for datasets like Thrombin and VEGF, whose experimental Ground Truth labels lack strong support from independent binding affinity measurements. Isolating the threshold selection from label reliance thus protects the classification from potential label errors. However, the approach still encounters label-independent limitations. In the ASSET dataset, the largest RCI gap splits two Active sequences rather than separating the Active and Inactive classes (which instead divide at the second-largest gap). Consequently, the rule misclassifies one sequence (ASSET_Active1) as Inactive, proving that the widest gap may not always reflect the true biological boundary—especially when a single class covers a broad RCI spread. Rather than tweaking the threshold *post hoc* and reintroducing label bias into an unlabelled routine, Table S2 explicitly flags this anomaly to maintain procedural integrity.

Several limitations should be acknowledged. Landscapes were derived from secondary structure models and did not explicitly incorporate tertiary interactions, ligand-induced folding, or kinetic pathways. Calculations assumed equilibrium thermodynamics under standard ionic conditions, and enrichment dynamics were analyzed retrospectively rather than modeled mechanistically. Nonetheless, the consistent association between elevated ruggedness and activity attrition across datasets suggests that intrinsic folding organization constrains activity resilience.

## CONCLUSION

This work identifies folding-landscape topology as a general structural characteristic that can be associated with aptamer activity. Our studies support a landscape-centric model of aptamer activity in which activity competence is associated not merely with thermodynamic stability but with the coherent multiscale organization of the folding ensemble. The RCI provides a compact, mechanistically grounded summary of this organization and offers a practical adjunct to enrichment-based prioritization in aptamer discovery. Framing aptamer activity through this lens may clarify observations from SELEX experiments in which some sequences show initial short-lived enrichment yet fail to maintain specificity. Such outcomes can be understood as consequences of disordered or competitive conformational ensembles, indicating that enrichment trends alone do not fully reflect viable activity.

In practice, the RCI may be seen as a measure that reflects ensemble organization directly from sequence. When used alongside conventional analyses, it offers a way to refine candidate selection early in the SELEX process.

## MATERIALS AND METHODS MATERIALS

### Dataset Compilation and Activity Definitions

Aptamer sequences were obtained from previously published studies spanning diverse molecular targets, experimental strategies, and validation criteria (Table S3). The six datasets analyzed were ASSET, IL_10, VEGF, Chloramphenicol, Dengue, and Thrombin.^14,35–39^ These datasets provided heterogeneous yet experimentally characterized systems for evaluating whether folding energy landscape organization can distinguish active from inactive aptamers across different biological contexts.

The ASSET dataset was derived from a high-throughput SELEX (HT-SELEX) experiment targeting vascular smooth muscle cells and contained 363 unique aptamer sequences, including 115 sequences classified as active and 248 classified as inactive based on experimentally determined specificity scores derived from next-generation sequencing analysis of target and non-target cell binding assays. Activity labels were assigned according to the enrichment and specificity criteria reported in the original ASSET study. The IL_10 dataset consisted of 42 RNA aptamer sequences selected against the IL_10 receptor (IL_10RA), for which activity classification was based on experimentally measured dissociation constants (Kd), with lower-affinity constants being classified as active and higher values being classified as inactive.

The remaining datasets were compiled from published experimental studies of well-characterized aptamers targeting distinct biomolecular classes. The VEGF dataset (n = 8 sequences) included DNA aptamers isolated through SELEX against vascular endothelial growth factor, with activity defined by reported binding affinity and retention in aptamer-binding assays. The Chloramphenicol dataset (n = 6 sequences) consisted of small-molecule-binding aptamers reported in biosensing and affinity-selection studies, in which activity was determined using reported binding affinity measurements or functional sensor performance described in the original publications. The Dengue dataset (n = 7 sequences) contained RNA aptamers selected against dengue viral proteins, with activity defined by experimentally validated binding to viral targets in biochemical assays. The Thrombin dataset (n = 15 sequences) included well-characterized thrombin-binding DNA aptamers whose activity was defined by reported binding affinity measurements and functional inhibition assays.

### Determination of Active Status and Short-Lived Enrichment

Active versus inactive classifications were defined based on published experimental benchmark data derived for each dataset. Briefly, for the primary datasets, activity was defined as follows:

ASSET HT-SELEX Dataset: Active sequences were defined using experimentally determined specificity scores calculated by normalized next-generation sequencing (NGS) across target vs. non-target cell binding assays. Sequences with specificity scores exceeding a 4-fold threshold and reaching statistical significance (Student’s *t*-test) were classified as active, whereas sequences failing to meet these criteria were classified as inactive.

IL_10 Dataset: Classification was based on experimentally measured dissociation constants (*K*_d_) determined by binding assays, where sequences with high binding affinity (low *K*_d_) were classified as active and non-binding or low-affinity sequences as inactive.

Transiently enriched sequences were identified using round-by-round HT-SELEX tracking data (ASSET and IL_10). These are sequences that demonstrated strong positive fold-enrichment early in selection (e.g., Round 4 relative to Round 3 in ASSET or Round 4 relative to Round 2 in IL_10), but lost representation in later rounds (e.g., Round 9 relative to Round 8 in ASSET and Round 5 to Round 4 in IL_10) and were ultimately classified as inactive based on post-SELEX specificity and affinity assays.

All sequences analyzed corresponded to final or post-selection candidates rather than intermediate SELEX libraries. Enrichment information was used retrospectively (in ASSET) to compare early- and late-round behavior but was not incorporated directly into folding computations. Folding landscape analysis was performed exclusively using intrinsic sequence-derived thermodynamic ensembles.

## METHODS

An overview of the proposed method is provided in Figure S12. In the following, we describe the key steps comprising the method.

### Secondary Structure Prediction and Ensemble Generation

Secondary structure ensembles were computed using the ViennaRNA package (v2.5.1).^41^ For each aptamer sequence, the MFE structure was determined, along with the partition function and corresponding base-pairing probabilities. In addition, suboptimal structures within an energy window of up to 7 kcal·mol⁻¹ above the MFE were generated. When insufficient suboptimal structures were enumerated to adequately sample the conformational space, Boltzmann-weighted sampling—typically generating 1000 structures—was performed to ensure effective representation of the structural ensemble. All folding calculations were performed under standard ViennaRNA thermodynamic parameters (37 °C, 1 M Na⁺), which approximated physiological ionic conditions in implicit solvent.

The resulting ensembles represent equilibrium secondary structure distributions and do not explicitly model tertiary contacts or kinetic folding pathways.

### Statistical characterization of energetic and structural landscape features

We performed detailed statistical analyses of energetic and structural landscape features on the ASSET dataset (363 aptamers) and the IL_10 dataset (42 aptamers). Other datasets had relatively small sample sizes, rendering them unsuitable for detailed statistical analysis. For both the ASSET and IL_10 datasets, classical thermodynamic descriptors as well as higher-order structural landscape features were extracted for each sequence to comprehensively characterize RNA secondary structure stability and conformational behavior. Thermodynamic measures included the minimum free energy (MFE), defined as the free energy of the most thermodynamically stable predicted secondary structure, and the energy gap, defined as the difference in free energy between the MFE structure and the next-lowest-energy structure, which provided an estimate of structural stability separation and fold specificity.

In addition to these thermodynamic quantities, ensemble diversity and the number of basins were calculated to capture properties not reflected by single-structure energy measures. Ensemble diversity quantifies the average structural dissimilarity among conformations within the thermodynamic ensemble and therefore reflects conformational flexibility. The number of basins, representing distinct local minima in the folding energy landscape, serves as a measure of landscape ruggedness and structural heterogeneity. These non-thermodynamic features were included because biological activity is influenced not only by absolute stability but also by conformational variability, alternative folding pathways, and landscape complexity—properties that cannot be fully described by scalar energy values alone.

Group-level comparisons between active and inactive sequences were performed using standard descriptive statistics, including the mean, median, and interquartile range, and two-sample t-tests were applied where appropriate. Distribution-level statistical comparisons were conducted for the ASSET and IL_10 datasets due to their large sample sizes and uniform activity labels. Other datasets contained relatively fewer sequences and heterogeneous activity definitions; therefore, formal statistical testing of scalar energetic features was not conducted for those sets. Sequence length and GC content were also determined to evaluate potential compositional confounding effects. Finally, nonlinear dimensionality reduction techniques, including t-SNE, UMAP, and MDS, were applied to visualize the structural feature space; these embeddings were used solely for qualitative inspection and did not influence subsequent quantitative analyses.

### Structural Distance Measure

Structural dissimilarity between secondary structures was quantified using base-pair distance, defined as the number of base pairs present in one structure but absent in the other. This measure captures elementary rearrangements in secondary structure space and provides a natural measure of conformational proximity.^42–44^

### Definition of Energy Level Sets

Let *S* denote the ensemble of secondary structures with associated free energies *G*(*s*). For a given energy threshold Δ*G*, the level set *L*(Δ*G*) is defined as:

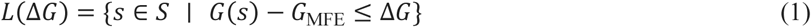

In Equation (1), *G*(*s*) denotes the Gibbs free energy (in kcal·mol⁻¹) of a given secondary structure *s*. Energies were computed using the ViennaRNA nearest-neighbor thermodynamic model, and all calculations were performed at 37°C under standard salt conditions (1 M Na⁺). Thus, *G*(*s*)corresponds to the predicted equilibrium free energy of that specific secondary structure.

The minimum free energy, *G*_MFE_, is defined as

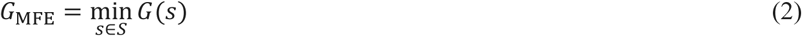

where *G*_MFE_ represents the lowest free energy among all structures in the ensemble The structure achieving this minimum is interpreted as the predicted thermodynamic ground state, and all *ΔG* calculations are defined relative to this reference energy. The lowest-energy structure (*ΔG* = 0, *i.e.,* the MFE structure) is considered first and then gradually structures with progressively higher free energy differences (*ΔG* > 0) are included in the analysis.

As the *ΔG* threshold is progressively increased, additional conformations are revealed that are less thermodynamically stable but still accessible within the ensemble. In other words, when *ΔG* = 0, only the single most stable structure is considered. As *ΔG* increases slightly (*e.g.,* 1–2 kcal·mol⁻¹), additional structures that are very close in energy to the ground state are included. At larger ΔG values (*e.g.,* 5–7 kcal·mol⁻¹), a substantially broader set of alternative folds becomes visible, reflecting increasingly diverse conformations within the ensemble. “Filtration” refers to this gradual inclusion of structures according to increasing energy thresholds. At each energy level, the relationships among structures can be examined—whether they form clusters, how they connect via small base-pair changes, and whether previously separate groups merge. The term “multiscale description” refers to analysis of the folding landscape across different energy scales. Low *ΔG* values highlight the most stable and tightly connected conformations, while higher *ΔG* values reveal the broader landscape, including alternative folds and larger-scale structural rearrangements.

Connected components of the graph at each Δ*G* define conformational basins. Basin size is measured as the number of structures within each component. As Δ*G* increases, new basins may emerge or existing basins may merge. Tracking these changes provides a dynamic description of fragmentation and landscape organization.

### Connectivity Graph Construction

For each level set *L*(Δ*G*), an undirected graph *G*_ΔG_ was constructed to represent structural connectivity within the energy threshold. In this graph, each node corresponds to a secondary structure *s* ∈ *L*(Δ*G*). An edge is placed between two nodes *s*_i_ and *s*_j_whenever their base-pair distance satisfies

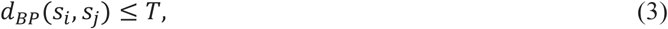

where *d*_BP_(*s*_i_, *s*_j_)denotes the base-pair distance—that is, the number of base pairs present in one structure but absent in the other—and *T*is a predefined distance threshold.

The parameter *T* controls the notion of structural adjacency and thus determines how permissive the connectivity criterion is. Small values of *T* restrict edges to structures differing by only minimal rearrangements, whereas larger values increasingly connect more distant conformations. In the context of RNA secondary structure space, small base-pair distances correspond to local structural modifications such as single base-pair shifts, limited helix fraying events, or minor stem adjustments. Consequently, imposing a distance threshold operationalizes an assumption of kinetic accessibility under local rearrangements.

To determine an appropriate value for *T*, sensitivity analyses were performed using nearby thresholds (e.g., *T* = 1, *T* = 2, and *T* = 3). When *T* = 1, the resulting graphs were highly fragmented, with the number of connected components increasing by approximately 30–45% relative to *T* = 2 across datasets, frequently separating structures that differed only by minor local rearrangements such as single base-pair shifts. In contrast, when *T* = 3, connectivity increased substantially, reducing the number of components by roughly 20–35% compared to *T* = 2 and occasionally merging clusters that remained structurally distinct under more conservative criteria. Across datasets, *T* = 2 consistently produced connectivity patterns that balanced fragmentation and over-merging while preserving the qualitative trends observed across energy levels. Because increasing or decreasing the threshold by one unit did not materially alter the overarching structural organization, the analysis was reliable to modest variations in *T*. Based on this empirical evaluation, *T* = 2 was selected as a principled compromise between structural resolution and graph coherence.

### *ΔG*-Dependent Topological Measures

To quantitatively capture how the RNA structural ensemble evolved across energy levels, three *ΔG*-dependent topological measures were computed for each sequence. These measures extended the level set framework by summarizing not just which structures are present below a given energy threshold, but also how they are organized and connected. First, the number of connected components in the connectivity graph of each level set measured the fragmentation of the ensemble, indicating whether structures form isolated basins or a largely connected network. Second, basin entropy, defined as the Shannon entropy of the normalized sizes of these connected components, quantified the dispersion of probability or structural weight among basins, thereby providing a measure of conformational diversity and accessibility. Finally, the cumulative structure count was used to track the total number of structures included at or below each *ΔG* threshold, reflecting how structural density grows as higher-energy conformations are progressively revealed. Together, these measures offered a multiscale, interpretable description of the folding landscape by characterizing how connectivity, basin distribution, and structural richness changed across energy levels, transforming the raw level sets into meaningful descriptors of RNA ensemble organization.

### The Ruggedness Composite Index (RCI)

The Ruggedness Composite Index (RCI) integrates four complementary features: the fragmentation of structures across ΔG levels, basin entropy reflecting the distribution of structural weight among connected components, variability in cumulative structure density growth (capturing slope irregularity), and the correlation length between structural distance and energy (ξ). The correlation length quantifies how rapidly free energy decorrelates with structural distance; shorter correlation lengths indicate sharper energetic fluctuations and increased local roughness. Together, these features combine into a single measure that characterizes the landscape’s connectivity, diversity, and energetic ruggedness, as described in Equation (4).

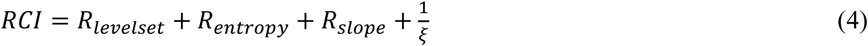

Each component captures a distinct aspect of folding organization: global fragmentation, entropic dispersion, energetic irregularity, and local smoothness. In order to assess potential redundancy among components, we performed pairwise regression analysis (Figure S13) on the ASSET aptamer dataset. Pairwise comparisons among level set fragmentation, basin entropy, cumulative density slope, and correlation length yielded low coefficients of determination and largely non-significant p-values, indicating weak linear dependence. This minimal redundancy supported the use of an additive RCI formulation without interaction terms. On this basis, an additive RCI formulation was adopted to preserve interpretability and avoid overfitting. Higher RCI values correspond to increasingly fragmented, high-entropy, and irregular landscapes, whereas lower values indicate smoother, funnel-like organization.

RCI-based activity thresholds, resubstitution and leave-one-out-cross validation procedures and full confusion-matrix statistics for all six aptamer datasets are provided in Tables S2, Table 1, Table 2

## Data Availability Statement

Raw data and additional information will be available upon request to the corresponding author.

## Acknowledgements

This work was supported in part by the National Science Foundation grants #IIS-1817239 (RS) and #IIS-3401096 (RS), Dr. Ralph and Marian Falk Medical Research Trust (RS, WT) and NHLBI R01HL157956 (WT)

## Author Contributions

G.S. (Gopalakrishnan Subramanian) conceived the computational framework, designed and implemented the energy landscape analysis pipeline, performed data analysis, and drafted the manuscript.

W.T. (William Thiel) contributed to biological interpretation of aptamer datasets, provided domain expertise on SELEX and experimental aptamer systems, and assisted in manuscript review and refinement.

R.S. (Rahul Singh) motivated the Level Set approach and supervised the study, guided methodological development and theoretical framing, contributed to interpretation of results, and critically revised the manuscript.

All authors reviewed, edited, and approved the final manuscript.

## Supplemental Information

Figures S1-S13, Tables S1-S3 and Supplemental Computational Implementation

## References

1. M. He, Z. Wang, X. Wu, Z. Du, C. Cui, Z. Zhao, Y. Sun, X. Zhang, L. He, W. Tan, (2025). Functional SELEX and Biomedical Applications of Aptamers: Beyond Molecular Recognition. Angew. Chem. Int. Ed., 64, e202424687. 10.1002/anie.202424687

2. Zhu C, Feng Z, Qin H, Chen L, Yan M, Li L, Qu F. (2024). Recent progress of SELEX methods for screening nucleic acid aptamers. Talanta. Jan 1;266(Pt 1):124998. doi: 10.1016/j.talanta.2023.124998. Epub 2023 Jul 26. PMID: 37527564.

3. Fang Z, Feng X, Tang F, Jiang H, Han S, Tao R, Lu C. (2024). Aptamer Screening: Current Methods and Future Trend towards Non-SELEX Approach. Biosensors (Basel). Jul 18;14(7):350. doi: 10.3390/bios14070350. PMID: 39056626; PMCID: PMC11274700.

4. Thiel WH, Bair T, Wyatt Thiel K, Dassie JP, Rockey WM, Howell CA, Liu XY, Dupuy AJ, Huang L, Owczarzy R, (2011) Nucleotide bias observed with a short SELEX RNA aptamer library. Nucleic Acid Ther.;21(4):253–63. Epub 2011/07/29. doi: 10.1089/nat.2011.0288. PubMed PMID: 21793789; PMCID: 3198618.

5. Takahashi M, Wu X, Ho M, Chomchan P, Rossi JJ, Burnett JC, Zhou J. (2016). High throughput sequencing analysis of RNA libraries reveals the influences of initial library and PCR methods on SELEX efficiency, Sci Rep.;6:33697. Epub 20160922. doi: 10.1038/srep33697. PubMed PMID: 27652575; PMCID: PMC5031971.

6. Wang J, Rudzinski JF, Gong Q, Soh HT, Atzberger PJ. (2012). Influence of Target Concentration and Background Binding on In Vitro Selection of Affinity Reagents. PLoS ONE. 7(8): e43940. doi: 10.1371/journal.pone.0043940.

7. Tolle F, Wilke J, Wengel J, Mayer G. (2014). By-Product Formation in Repetitive PCR Amplification of DNA Libraries during SELEX. PLoS ONE. 9(12):e114693. doi: 10.1371/journal.pone.0114693.

8. Alkhamis O, Xiao Y. (2023). Systematic Study of in Vitro Selection Stringency Reveals How To Enrich High-Affinity Aptamers. Journal of the American Chemical Society. 145(1):194–206. doi:10.1021/jacs.2c09522.

9. Paolo Climaco, Noelle M. Mitchell, Matthew J. Tyler, Kyungae Yang, Anne M. Andrews, Andrea L. Bertozzi, (2025) GMFOLD: Subgraph matching for high-throughput DNA-aptamer secondary structure classification and machine learning interpretability, Mathematical Biosciences, Volume 387, 109485, ISSN 0025-5564.

10. Chen, L., Zhang, B., Wu, Z., Liu, G., Li, W. and Tang, Y., (2023). In Silico discovery of aptamers with an enhanced library design strategy. Computational and Structural Biotechnology Journal, 21, pp.1005–1013.

11. Zhen Wang, Ziqi Liu, Wei Zhang, Yanjun Li, Yizhen Feng, Shaokang Lv, Han Diao, Zhaofeng Luo, Pengju Yan, Min He, Xiaolin Li, (2024). AptaDiff: de novo design andoptimization of aptamers based on diffusion models, Briefings in Bioinformatics, Volume 25, Issue 6, November Bbae 517. doi.org/10.1093/bib/bbae517

12. Schroeder SJ. (2018) Challenges and approaches to predicting RNA with multiple functional structures. RNA. (2018) Dec;24(12):1615–1624. doi: 10.1261/rna.067827.118. Epub Aug 24. PMID: 30143552; PMCID: PMC6239171.

13. Qu H, Ma Q, Wang L, Mao Y, Eisenstein M, Soh HT, Zheng L. (2020). Measuring Aptamer Folding Energy Using a Molecular Clamp. J Am Chem Soc. Jul 8;142(27):11743–11749. doi: 10.1021/jacs.0c01570. Epub Jun 22. PMID: 32491843.

14. Rogers E, Heitsch C. (2016). New insights from cluster analysis methods for RNA secondary structure prediction. Wiley Interdiscip Rev RNA. May;7(3):278–94. doi: 10.1002/wrna.1334. Epub Mar 11. PMID: 26971529; PMCID: PMC4840050.

15. Jiang W, Jones JC, Shankavaram U, Sproull M, Camphausen K, Krauze AV. (2022). Analytical Considerations of Large-Scale Aptamer-Based Datasets for Translational Applications. Cancers (Basel). Apr 29;14(9):2227. doi: 10.3390/cancers14092227.35565358; PMCID: PMC9105298.

16. Jing M, Bowser MT. (2013). Tracking the emergence of high affinity aptamers for rhVEGF165 during capillary electrophoresis-systematic evolution of ligands by exponential enrichment using high throughput sequencing. Anal Chem. Nov 19;85(22):10761–70. doi: 10.1021/ac401875h. Epub Nov 1. PMID: 24125636; PMCID: PMC3892959.

17. Levay A, Brenneman R, Hoinka J, Sant D, Cardone M, Trinchieri G, Przytycka TM, Berezhnoy A. (2015). Identifying high-affinity aptamer ligands with defined cross-reactivity using high-throughput guided systematic evolution of ligands by exponential enrichment. Nucleic Acids Res. Jul 13;43(12): e82. doi: 10.1093/nar/gkv534. Epub May 24. PMID: 26007661; PMCID: PMC4499151.

18. Yan S, Ilgu M, Nilsen-Hamilton M, Lamm MH. (2022). Computational Modeling of RNA Aptamers: Structure Prediction of the Apo State. J Phys Chem B. Sep 22;126(37):7114–7125. doi: 10.1021/acs.jpcb.2c04649. Epub Sep 12. PMID: 36097649; PMCID: PMC9512008.

19. Savinov A, Perez CF, Block SM. (2014). Single-molecule studies of riboswitch folding. Biochim Biophys Acta. Oct;1839(10):1030–1045. doi: 10.1016/j.bbagrm.2014.04.005. Epub Apr 13. PMID: 24727093; PMCID: PMC4177941.

20. Lin JC, Hyeon C, Thirumalai D. (2014). Sequence-dependent folding landscapes of adenine riboswitch aptamers. Phys Chem Chem Phys. Apr 14;16(14):6376–82. doi: 10.1039/c3cp53932f. Epub Dec 23. PMID: 24366448; PMCID: PMC5580260.

21. R. Micura and C. Höbartner, (2020).“Fundamental studies of functional nucleic acids: aptamers, riboswitches, ribozymes and DNAzymes,” Chem. Soc. Rev. 49, 7331–7353 doi: 10.1039/D0CS00617C

22. Sato, K., & Hamada, M. (2023). Recent trends in RNA informatics: a review of machine learning and deep learning for RNA secondary structure prediction and RNA drug discovery. Briefings in Bioinformatics, 24(4), bbad186. 10.1093/bib/bbad186

23. Zuker M. (1989). On finding all suboptimal foldings of an RNA molecule. Science. Apr 7;244(4900):48–52. doi: 10.1126/science.2468181. PMID: 2468181.

24. McCaskill JS. (1990). The equilibrium partition function and base pair binding probabilities for RNA Secondary structure. Biopolymers. May-Jun;29(6-7):1105–19. doi: 10.1002/bip.360290621. PMID: 1695107.

25. Ding Y, Chan CY, Lawrence CE. (2004). Sfold web server for statistical folding and rational design of nucleic acids. Nucleic Acids Res. Jul 1;32(Web Server issue): W135–41. doi: 10.1093/nar/gkh449. PMID: 15215366; PMCID: PMC441587.

26. Onuchic, J. N., Luthey-Schulten, Z. & Wolynes, P. G. (1997). Theory of protein folding: the energy landscape perspective. Annu. Rev. Phys. Chem. 48, 545–600.

27. Ditzler MA, Rueda D, Mo J, Håkansson K, Walter NG. (2008). A rugged free energy landscape separates multiple functional RNA folds throughout denaturation. Nucleic Acids Res.

28. Volkhardt A, Grubmüller H. (2022). Estimating ruggedness of free-energy landscapes of small globular proteins from principal component analysis of molecular dynamics trajectories. Phys Rev E. Apr;105(4-1):044404. doi:10.1103/PhysRevE.105.044404. PMID 35590540.

29. Woodside MT, Block SM. (2014). Reconstructing folding energy landscapes by single-molecule force spectroscopy. Annu Rev Biophys.;43:19–39. doi: 10.1146/annurev-biophys-051013-022754. PMID: 24895850; PMCID: PMC4609573.

30. Solomatin SV, Greenfeld M, Chu S, Herschlag D. (2010). Multiple native states reveal persistent ruggedness of an RNA folding landscape. Nature. Feb 4;463(7281):681–4. doi: 10.1038/nature08717. PMID: 20130651; PMCID: PMC2818749.

31. Wales, D. J. (2003). Energy Landscapes: Applications to Clusters, Biomolecules and Glasses. Cambridge University Press.

32 Flamm, C., Fontana, W., Hofacker, I. L., & Schuster, P. (2000). RNA folding at elementary step resolution. RNA, 6(3), 325–338.

33. Krivov, S. V., & Karplus, M. (2004). Hidden complexity of free energy surfaces for peptide (protein) folding. Proceedings of the National Academy of Sciences, 101(41),14766–14770.

34. Kucharík, M., Hofacker, I. L., Stadler, P. F., & Flamm, C. (2014). An efficient algorithm for the exhaustive analysis of RNA energy landscapes. Bioinformatics, 30(11), 1697, 1703.

35. Barros M, Kasirajan G, Jones A, Schlichting A, Ruiz-Ciancio D, Lin LH, Narayan C, Veeramani S, Thiel KW, Kennedy GC, Darcy I, Thiel W. (2025). ASSET: A Framework for Decoding Aptamer Specificity of an Enriched Library by Next-Generation Sequencing of Experimental Samples. bioRxiv [Preprint]. Aug 30:.08.27.672406. doi: 10.1101/2025.08.27.672406. PMID: 40909745; PMCID: PMC12407934.

36. Kaur H, Yung LY. (2012). Probing high affinity sequences of DNA aptamer against VEGF165. PLoS One.;7(2):e31196. doi: 10.1371/journal.pone.0031196. Epub 2012 Feb 16. PMID: 22359573; PMCID: PMC3281051.

37. Burke DH, Hoffman DC, Brown A, Hansen M, Pardi A, Gold L (1997). RNA aptamers to the peptidyl transferase inhibitor chloramphenicol. Chem Biol. 1 Nov;4(11):833–43. doi: 10.1016/s1074-5521(97)90116-2. PMID: 9384530.

38. Thevendran R, Rogini S, Leighton G, Mutombwera A, Shigdar S, Tang TH, Citartan M. (2023). The Diagnostic Potential of RNA Aptamers against the NS1 Protein of Dengue Virus Serotype 2. Biology (Basel). May 15;12(5):722. doi: 10.3390/biology12050722. PMID: 37237536; PMCID: PMC10215423.

39. Kubik MF, Stephens AW, Schneider D, Marlar RA, Tasset D. (1994). High-affinity RNA ligands to human alpha-thrombin. Nucleic Acids Res. Jul 11;22(13):2619–26. doi: 10.1093/nar/22.13.2619. PMID: 7518917; PMCID: PMC308218

40. Woodson SA. (2010). Compact intermediates in RNA folding. Annu Rev Biophys.;39:61–77. doi: 10.1146/annurev.biophys.093008.131334. PMID: 20192764; PMCID: PMC6341483.

41. Ronny Lorenz, Stephan H. Bernhart, Christian Hönerzu Siederdissen, Hakim Tafer, Christoph Flamm, Peter F. Stadler, and Ivo L. Hofacker. (2011). “ViennaRNA Package 2.0”. Algorithms for Molecular Biology, 6(1):26. (DOI): 10.1186/1748-7188-6-26.

42. Mathews, D. H., & Turner, D. H. (2006). Prediction of RNA secondary structure by free energy minimization. Current Opinion in Structural Biology, 16(3), 270–278.

43. Chakraborty D, Collepardo-Guevara R, Wales DJ. (2014). Energy landscapes, folding mechanisms, and kinetics of RNA tetraloop hairpins. J Am Chem Soc. 2014 Dec 31;136(52):18052–61. doi: 10.1021/ja5100756. Epub Dec 17. PMID: 25453221.

44. Zhao, Y., Gao, B., Chen, Y. & Liu, J. (2023). An aptamer array for discriminating tetracycline antibiotics based on binding-enhanced intrinsic fluorescence. Analyst 148, 1507.

