## Supplemental Information for "A Simple Method to Distinguish Active and Inactive Aptamers by Analyzing the Ruggedness of the Aptamer Free Energy Landscape"

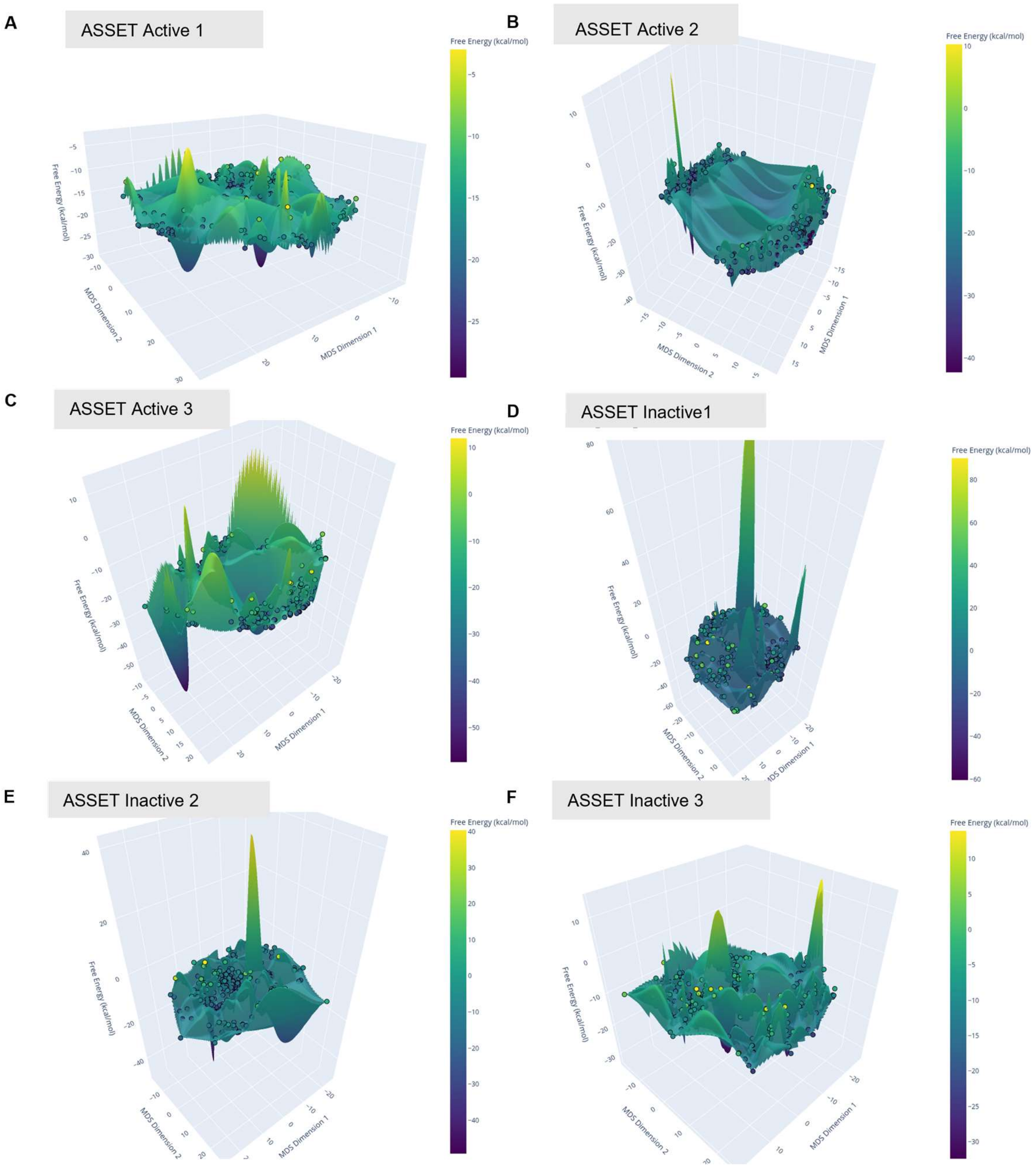


**Figure S1.** 3D Visualizations of Energy Landscapes of ASSET Active Aptamer1(A), ASSET Active Aptamer2(B), ASSET Active Aptamer3(C), ASSET Inactive Aptamer1(D), ASSET Inactive Aptamer2(E), ASSET Inactive Aptamer3(F),

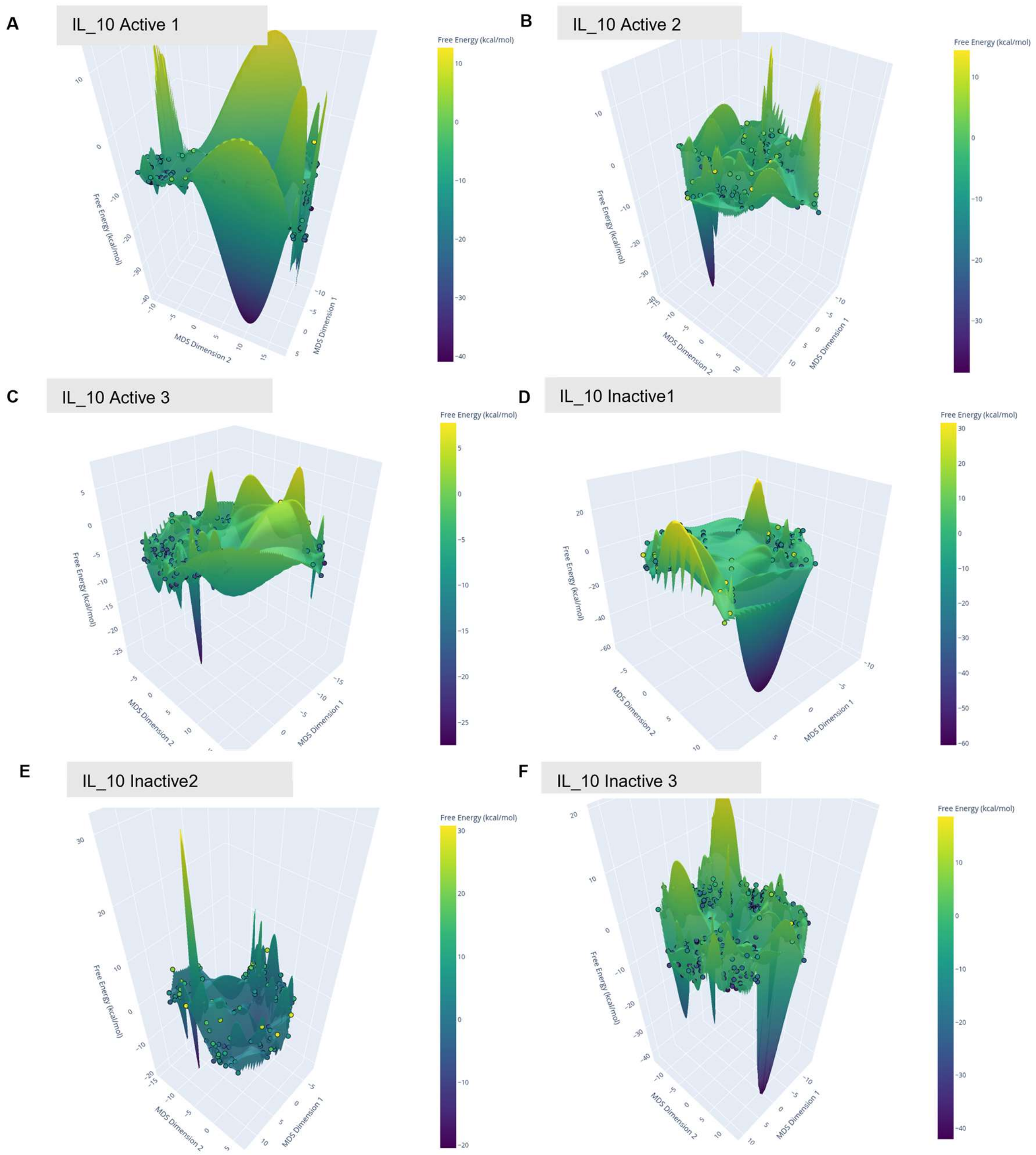


**Figure S2.** 3D Visualizations of Energy Landscapes of IL_10 Active Aptamer1(A), IL_!0 Active Aptamer2(B), IL_10 Active Aptamer3(C), IL_10 Inactive Aptamer1(D), IL_10 Inactive Aptamer2(E), IL_10 Inactive Aptamer3(F),

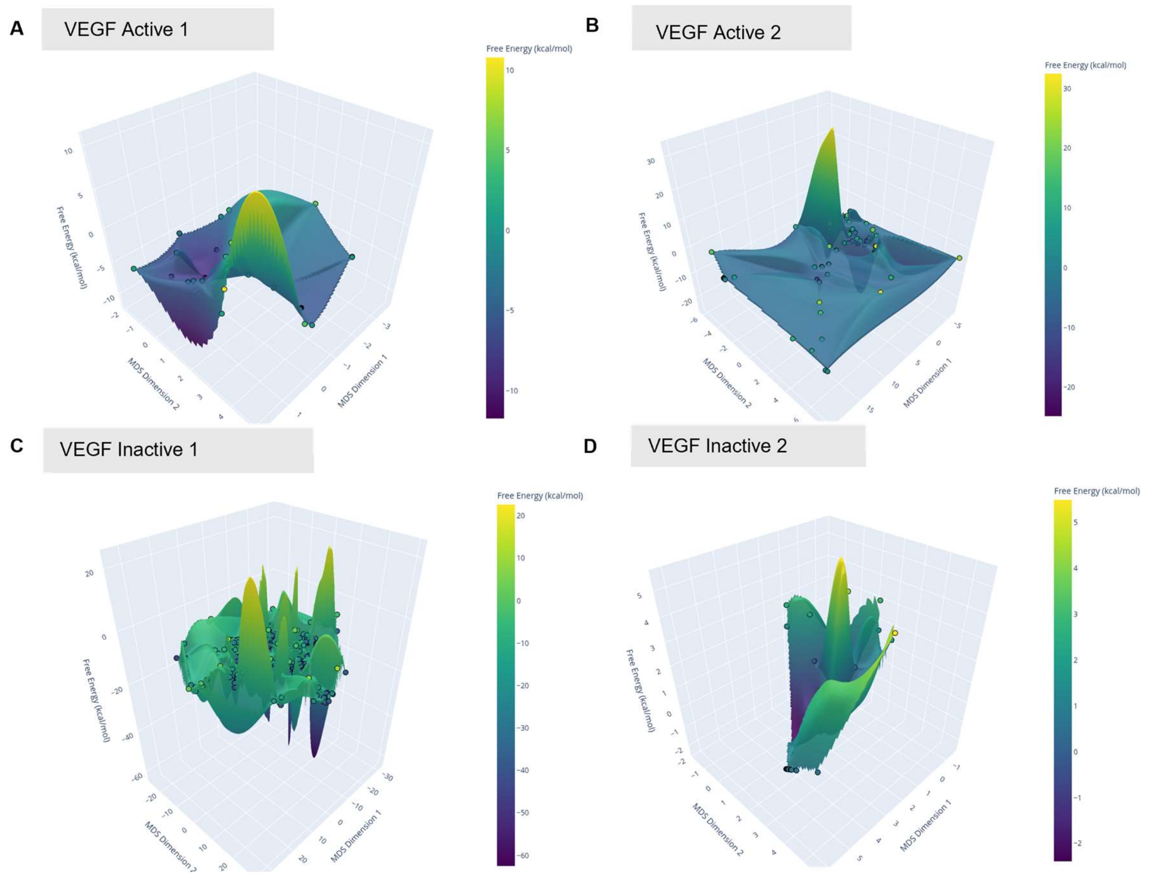


**Figure S3.** 3D Visualizations of Energy Landscapes of VEGF SL2B_Active1Aptamer (A), VEGF SL12_Active2 Aptamer (B), VEGF SL1_Inactive1Aptamer (C), VEGF VEa5_Inactive2 Aptamer (D)

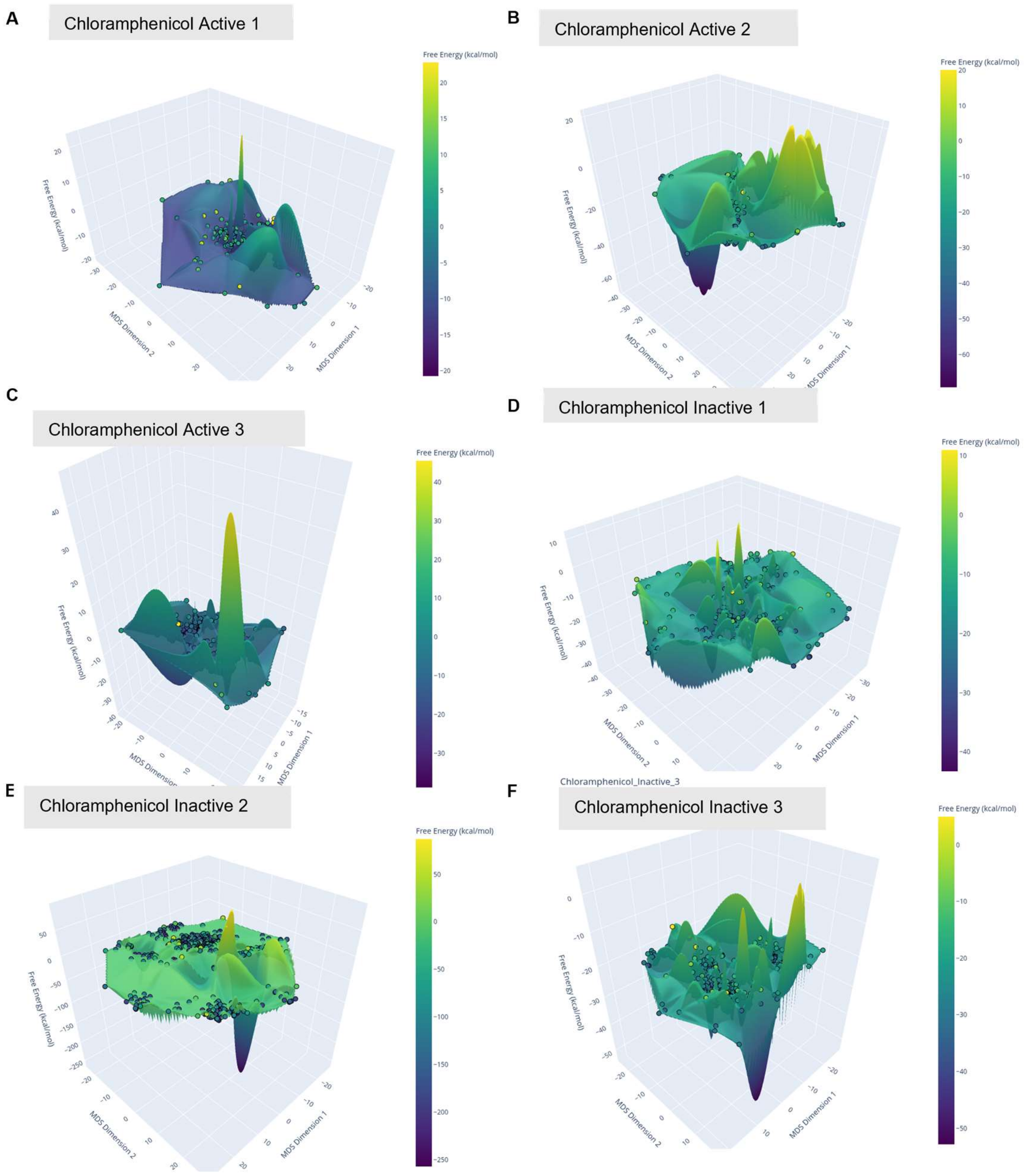


**Figure S4.** 3D Visualizations of Energy Landscapes of Chloramphenicol 70Cm6_ Active1 Aptamer (A), Chloramphenicol 70Cm9_Active2 Aptamer (B), Chloramphenicol 70Cm53_Active3 Aptamer (C) Chloramphenicol 80Cm51_ Inactive1 Aptamer (D), Chloramphenicol 80Cm38_Inactive2 Aptamer (E) and Chloramphenicol 80Cm54_Inactive3 Aptamer (F)

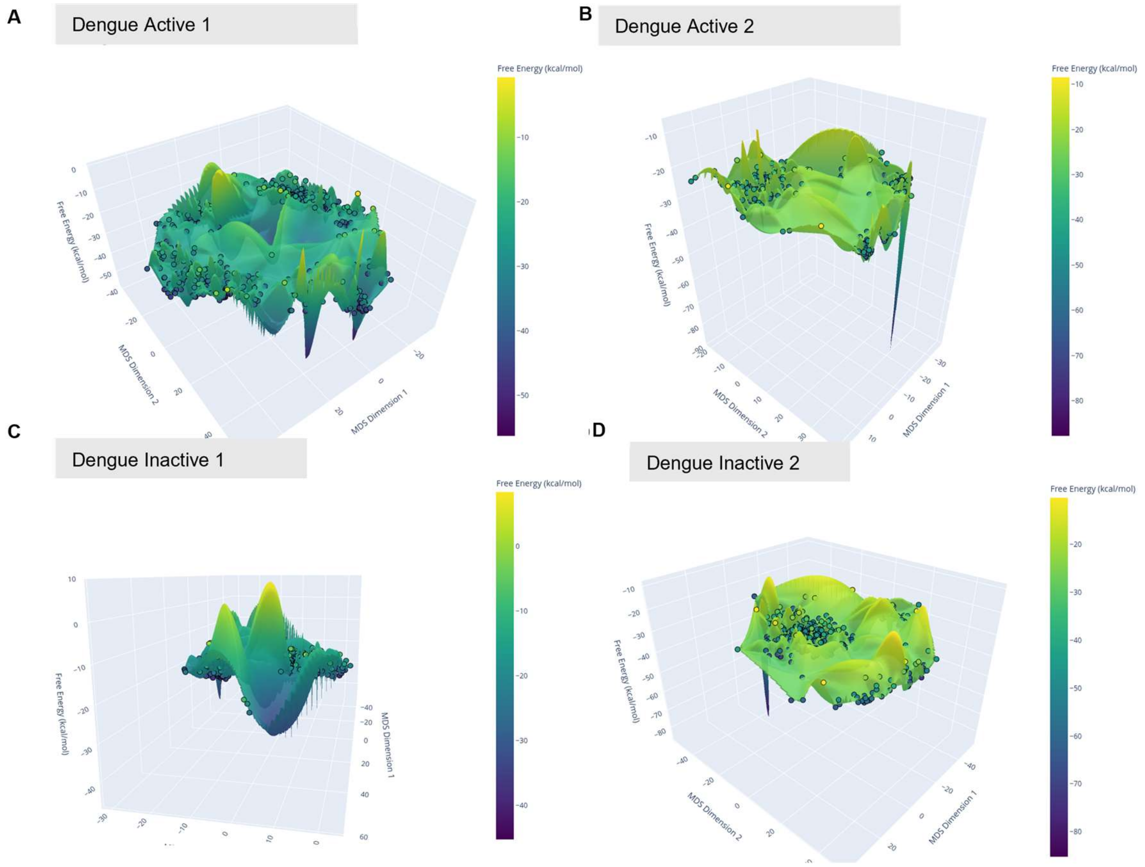


**Figure S5.** 3D Visualizations of Energy Landscapes of Dengue DENV-3_Active1 Aptamer (A), Dengue DENV-6_Active2 Aptamer (B), Dengue DENV-2_Inactive1Aptamer (C), Dengue DENV-7_Inactive2 Aptamer (D)

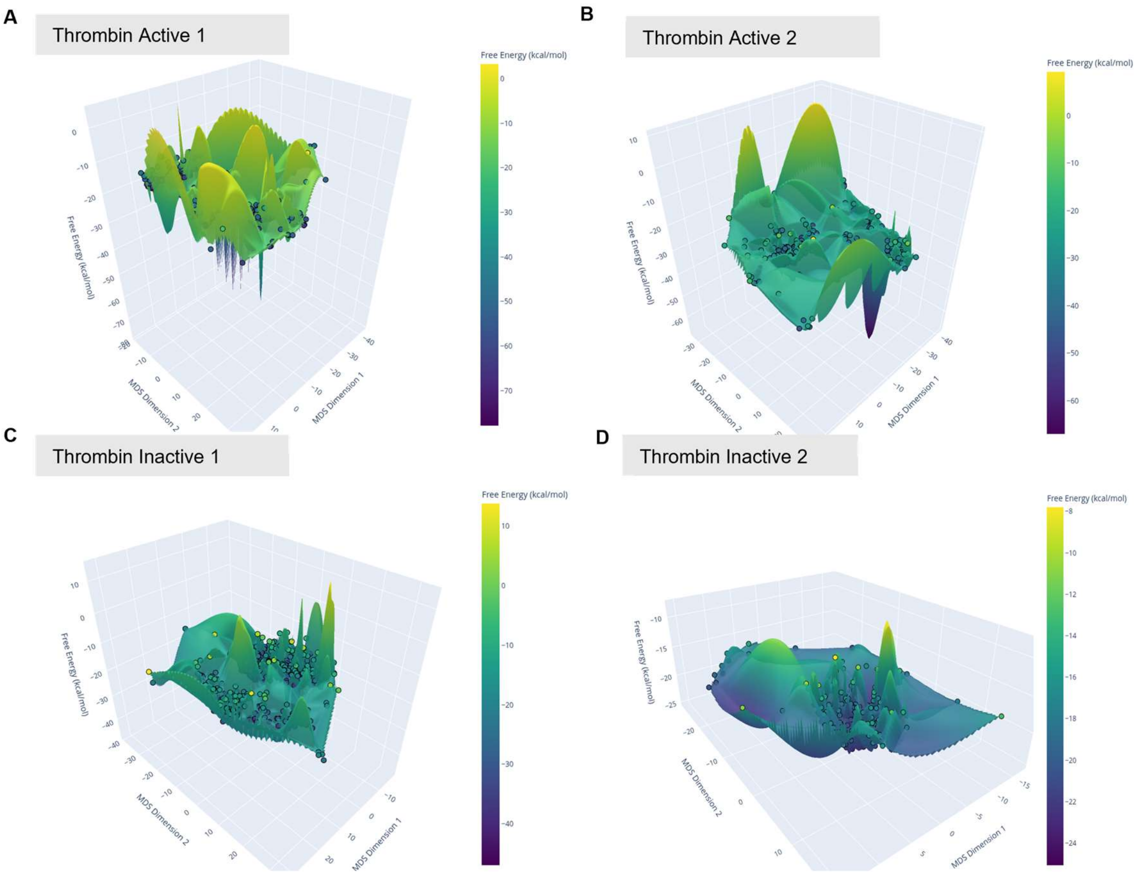


**Figure S6.** 3D Visualizations of Energy Landscapes of Thrombin 16.24 Active1 Aptamer (A), Thrombin 27.33 Active2 Aptamer (B), Thrombin 2_Inactive1 Aptamer (C), Thrombin 5_Inactive2 Aptamer (D)

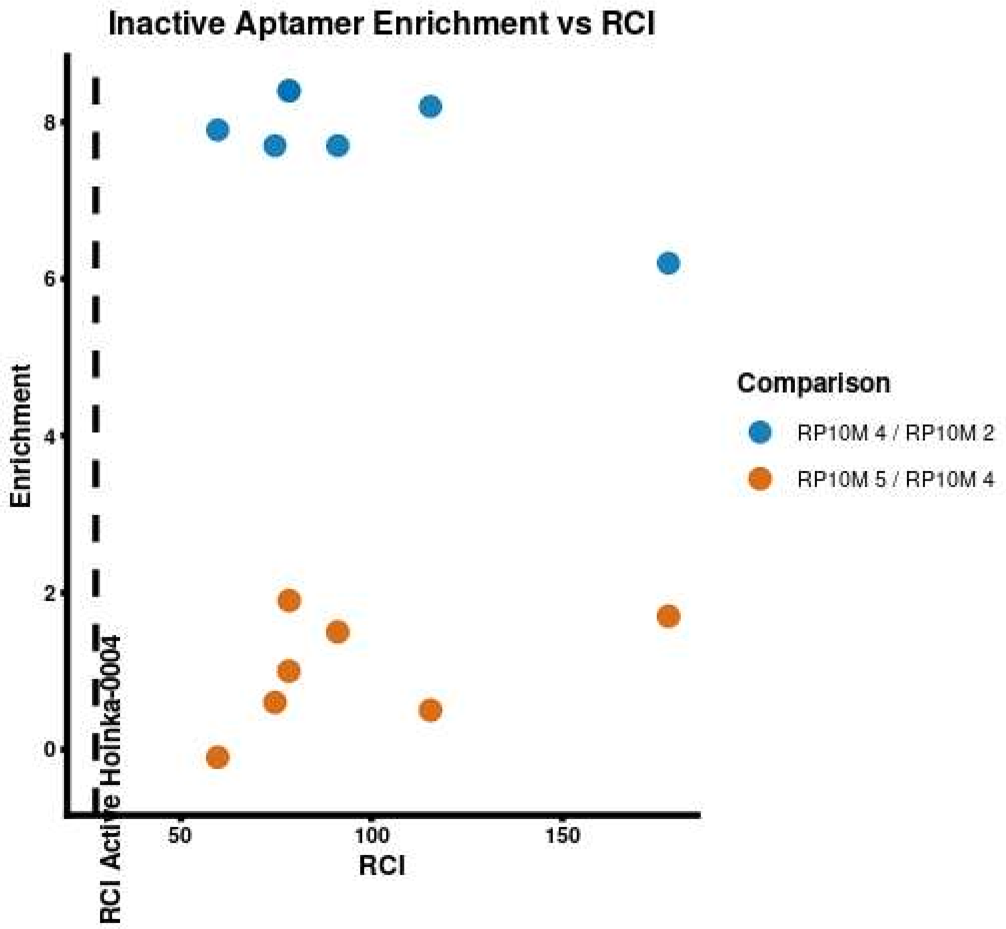


**Figure S7 Round-to-round enrichment behavior and folding landscape Ruggedness Composite Index (RCI) of Inactive IL 10 aptamers**. The figure shows sequence-level enrichment ratios across early and late SELEX rounds (Blue Points represent Round 4/ Round 2 and Orange Points represent Round 5/Read Round 4)) alongside the RCI computed from sequence-derived folding energy landscapes with low RCI of Active Hoinka-0004 aptamer as reference. All sequences exhibit significant positive enrichment at early rounds yet fail to retain enrichment at later rounds and are ultimately classified as inactive based on final Kd values. These early-enriched but inactive sequences consistently display elevated RCI values, whereas sequences that remain enriched through later rounds that are classified as active exhibit lower RCI values. The figure illustrates that landscape RCI provides an independent discriminator of functional outcome beyond enrichment trajectories alone

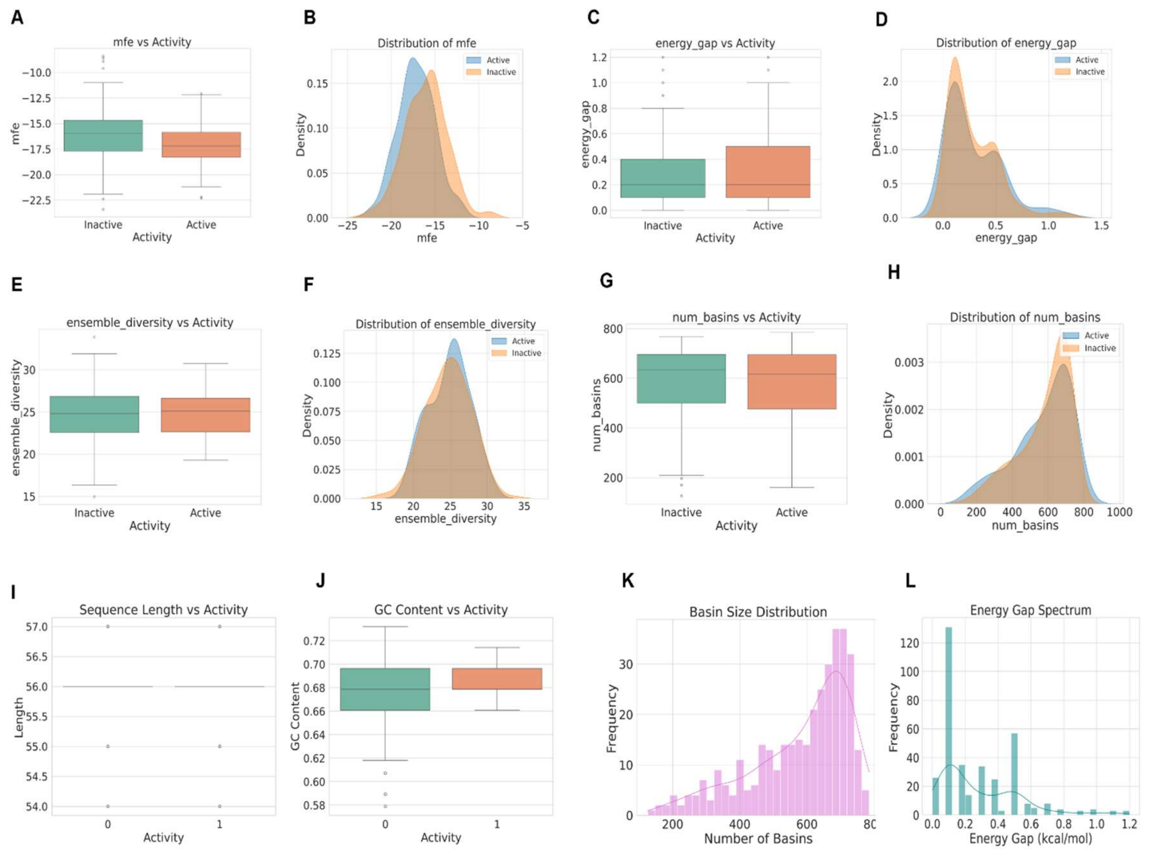


**Figure S8. Comparative analysis of structural and thermodynamic features distinguishing active and inactive ASSET aptamers.** Box plots and corresponding distributions are shown for minimum free energy (A–B), energy gap (C–D), ensemble diversity (E–F), and number of basins (G–H). Sequence-level characteristics, including length (I) and GC content (J), are also compared by activity. Landscape-level analyses (K–L) depict basin size distributions and the energy gap spectrum, highlighting that ASSET aptamer folding is characterized by multiple competing conformations rather than a single dominant minimum

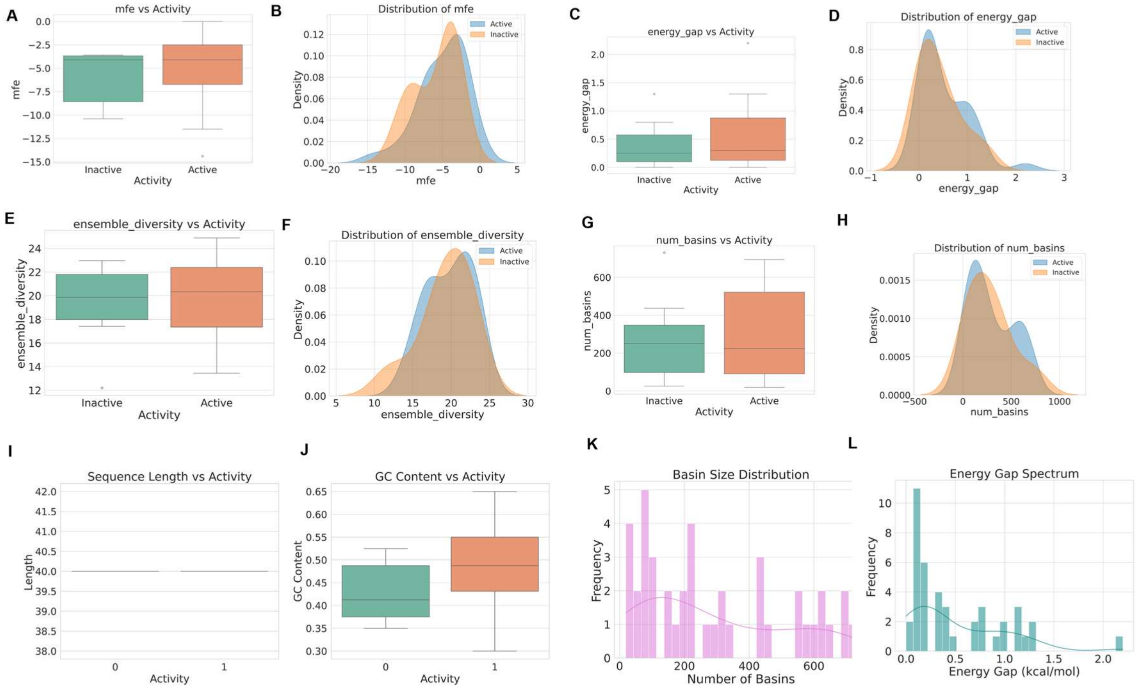


**Figure S9. Comparative analysis of structural and thermodynamic features distinguishing active and inactive IL-10 aptamers.** Box plots and corresponding distributions are shown for minimum free energy (A–B), energy gap (C–D), ensemble diversity (E–F), and number of basins (G–H). Sequence-level characteristics, including length (I) and GC content (J) are also compared by activity. Landscape-level analyses (K–L) depict basin size distributions and the energy gap spectrum, highlighting that IL-10 aptamer folding is characterized by multiple competing conformations rather than a single dominant minimum

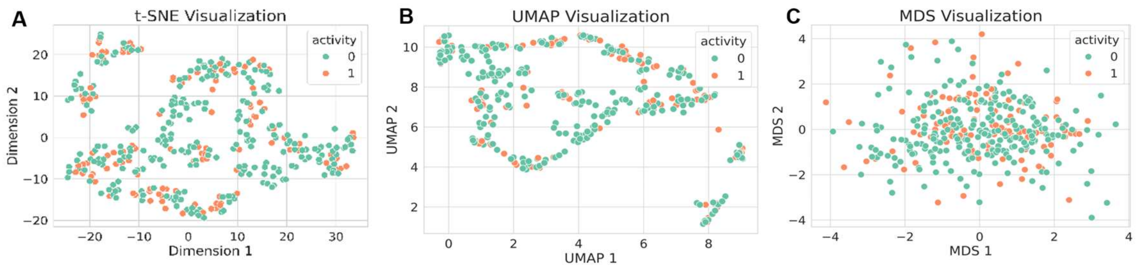


**Figure S10.** t-SNE (A), UMAP (B), and MDS (C) visualizations of structural clustering of active and inactive ASSET Aptamers

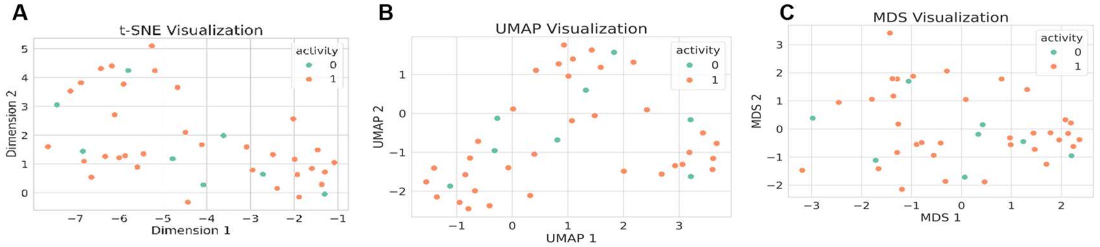


**Figure S11**. t-SNE (A), UMAP (B), and MDS (C) visualizations of structural clustering of active and inactive IL 10 aptamers.

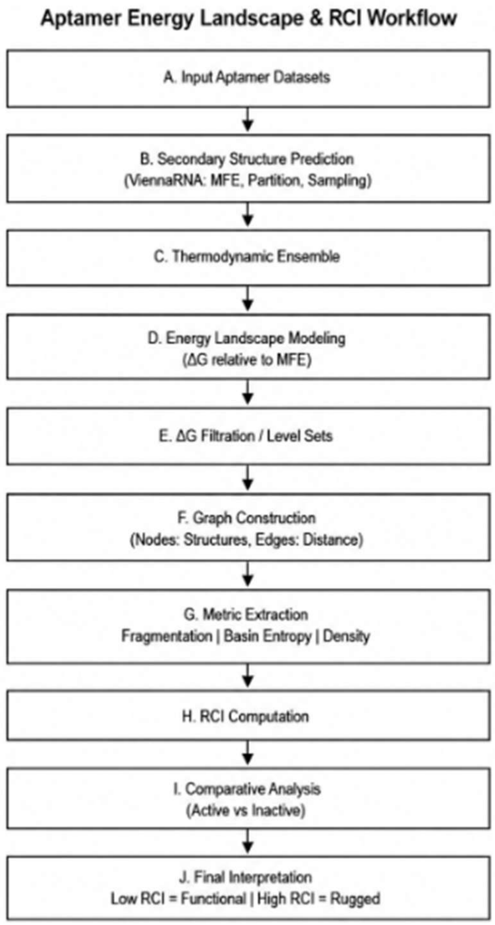


**Figure S12. Schematic of the computational pipeline for RCI-based analysis of aptamer energy landscapes.** Secondary structure ensembles generated by ViennaRNA are used to construct *ΔG*-dependent energy landscapes and connectivity graphs. Structural measures (fragmentation, basin entropy, and density) are computed across energy levels and integrated into the Ruggedness Complexity Index (RCI), enabling discrimination between active (low RCI) and inactive (high RCI) aptamers.

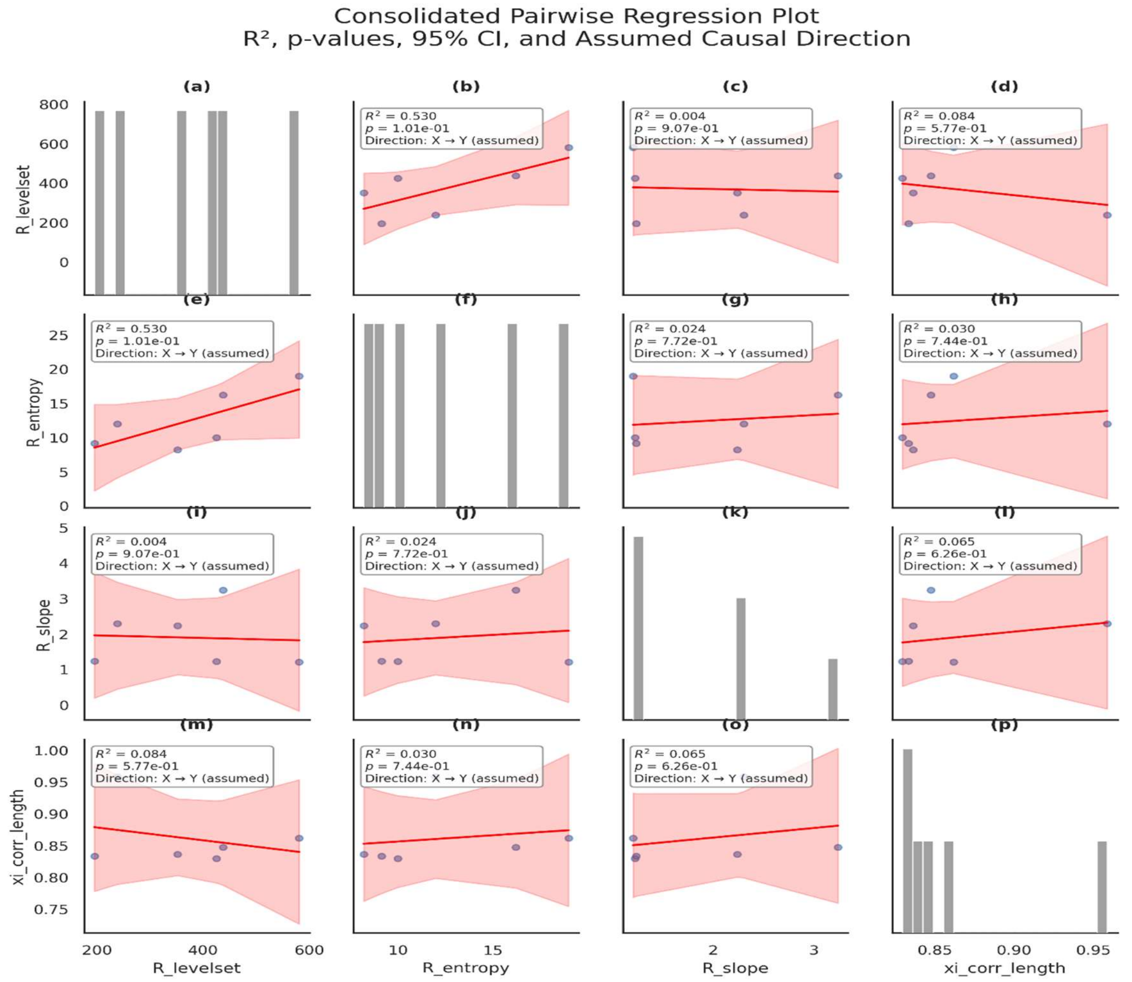


**Figure S13. Pairwise regression analysis among RCI components.** Each panel shows pairwise linear regressions between candidate RCI components (R_levelset, R_entropy, R_slope, and ξ-correlation length), with shaded regions indicating 95% confidence intervals. Reported values include coefficient of determination ($R^2$) and associated *p*-values. Diagonal panels show marginal histograms of each component of RCI, where bar heights represent the number of observations falling within each value range, providing context for sample size, spread, and distributional structure underlying the pairwise regressions. Across all pairwise comparisons, low $R^2$ values and non-significant *p*-values indicate weak linear dependence and minimal redundancy among components. Thus, these results support the use of an additive RCI formulation without explicit interaction terms, as each component captures largely independent aspects of the folding energy landscape

**Table S1**. Experimentally characterized aptamer datasets used for energy landscape analysis. The table summarizes the molecular targets, data sources, sequence counts, and activity definitions for the six aptamer datasets analyzed in this work. All datasets were compiled from prior publications and used without modification for comparative folding energy landscape analysis

| Dataset | Target | Origin / Selection Strategy | Number of Sequences | Activity Definition Used Here |
|---|---|---|---|---|
| ASSET Barros *et al.*, 2025[1] | Cell-surface / molecular targets | NGS-based specificity profiling of SELEX-derived libraries | 363 (115 Active, 248 Inactive) | Experimentally defined specificity classification |
| IL-10 Levay *et al*, 2015[2] | IL-10 receptor (IL-10RA) | SELEX with affinity-based optimization | 42 | Low Kd = Active; High Kd = Inactive |
| VEGF Kaur & Yung, 2012[3] | VEGF165 protein | Structure-guided truncation of known aptamer | 5 | Retained binding vs loss of binding |
| Chloramphenicol Burke *et al.,* 1997[4] | Chloramphenicol (small molecule) | Classical in vitro SELEX | 6 | Validated binding vs non-binding |
| Dengue Thevendran *et al*., 2023[5] | Dengue virus NS1 protein | In vitro SELEX against viral antigen | 4 | Experimental binding validation |
| Thrombin Kubik *et al*., 1994[6] | Human α-thrombin | Classical SELEX from | 4 | Reported high vs low affinity |

combinatorial

RNA pool

**Table S2. Sequence-Level Aptamer Classification: Ruggedness Composite Index (RCI) Predictions vs. Experimental Ground Truth.** This dataset compiles individual aptamers from six source files underlying Figures 3–5, pairing each sequence's calculated RCI with its experimental Ground Truth activity class and its RCI-derived Predicted activity class. The compilation includes entries for VEGF ($n = 4$), Dengue ($n = 4$), Thrombin ($n = 4$), ASSET ($n = 6$), and IL-10 ($n = 6$)—which generate the groupwise RCI statistics in Figures 3–5—alongside the full dataset for Chloramphenicol ($n = 6$). The decision rule assigns an **Active** label when an aptamer's RCI falls below a dataset-specific threshold (stated in the row preceding each dataset) and an **Inactive** label otherwise. To calculate each threshold independently of Ground Truth labels, the protocol sorts all RCI values in a dataset, identifies the sparsest interval between RCI scores distributions, and sets the cutoff at that interval's midpoint; rows with misaligned predictions appear in bold. For Chloramphenicol, the largest gap aligns perfectly with the true Active/Inactive boundary to yield clean separation. For VEGF, Dengue, Thrombin, and IL-10, the largest gap falls near the true boundary, misclassifying one or two sequences based on their position relative to the threshold rather than their true group label. For ASSET, the primary gap lies within the Active group itself, while the actual boundary gap ranks second largest. Specific caveats for each dataset appear in the header row preceding its entries.

| Aptamer ID (Label) | Data Set | RCI | Ground Truth | Predicted | Correct |
|---|---|---|---|---|---|
| ***VEGF (n = 4) RCI threshold = 231.90, the midpoint of the largest gap (sparsest interval) in the dataset's full sorted RCI distribution. largest gap (342.82) falls between VEGF_Active2 (60.49) and VEGF_Inactive2 (403.31); VEGF_Inactive1 (20.58) falls below the gap despite its Inactive label.*** | | | | | |
| VEGF_Active1 | VEGF | 16.27 | Active | Active | Yes |
| VEGF_Active2 | VEGF | 60.49 | Active | Active | Yes |
| VEGF_Inactive1 | VEGF | 20.58 | Inactive | **Active** | **No** |
| VEGF_Inactive2 | VEGF | 403.31 | Inactive | Inactive | Yes |

| Aptamer ID (Label) | Data Set | RCI | Ground Truth | Predicted | Correct |
|---|---|---|---|---|---|
| ***Chloramphenicol (n = 6) RCI threshold = 274.21, the midpoint of the largest gap (sparsest interval) in the dataset's full sorted RCI distribution. largest gap (157.98) falls exactly at the Active/Inactive group boundary, between Chloramphenicol_Active2 (195.22) and Chloramphenicol_Inactive1 (353.20) - clean separation.*** | | | | | |
| Chloramphenicol_Active1 | Chloramphenicol | 150.85 | Active | Active | Yes |
| Chloramphenicol_Active2 | Chloramphenicol | 195.22 | Active | Active | Yes |
| Chloramphenicol_Active3 | Chloramphenicol | 172.20 | Active | Active | Yes |
| Chloramphenicol_Inactive1 | Chloramphenicol | 353.20 | Inactive | Inactive | Yes |
| Chloramphenicol_Inactive2 | Chloramphenicol | 465.63 | Inactive | Inactive | Yes |
| Chloramphenicol_Inactive3 | Chloramphenicol | 377.26 | Inactive | Inactive | Yes |
| ***Dengue (n = 4) RCI threshold = 696.57, the midpoint of the largest gap (sparsest interval) in the dataset's full sorted RCI distribution. largest gap (638.64) falls between Dengue_Active2 (377.25) and Dengue_Inactive2 (1015.89); the high-RCI outlier Dengue_Active1 (1312.98) falls above the gap despite its Active label.*** | | | | | |
| Dengue_Active1 | Dengue | 1312.98 | Active | **Inactive** | **No** |
| Dengue_Active2 | Dengue | 377.25 | Active | Active | Yes |
| Dengue_Inactive1 | Dengue | 1065.12 | Inactive | Inactive | Yes |
| Dengue_Inactive2 | Dengue | 1015.89 | Inactive | Inactive | Yes |
| ***Thrombin (n = 4) RCI threshold = 628.05, the midpoint of the largest gap (sparsest interval) in the dataset's full sorted RCI distribution. largest gap (465.85) falls between Thrombin_Active1 (395.12) and Thrombin_Inactive1 (860.97); the low-RCI outlier Thrombin_Inactive2 (78.60) falls below the gap despite its Inactive label.*** | | | | | |
| Thrombin_Active1 | Thrombin | 395.12 | Active | Active | Yes |
| Thrombin_Active2 | Thrombin | 274.28 | Active | Active | Yes |
| Thrombin_Inactive1 | Thrombin | 860.97 | Inactive | Inactive | Yes |
| Thrombin_Inactive2 | Thrombin | 78.60 | Inactive | **Active** | **No** |
| ***ASSET (n = 6) RCI threshold = 294.05, the midpoint of the largest gap (sparsest interval) in the dataset's full sorted RCI distribution. largest gap (149.18) falls within the Active group itself, between ASSET_Active3 (219.46) and ASSET_Active1 (368.63), not at the Active/Inactive boundary; ASSET_Active1 therefore falls above the gap despite its Active label. The Active/Inactive boundary gap (102.81, between ASSET_Active1 and ASSET_Inactive2) is the second-largest gap in this dataset.*** | | | | | |
| ASSET_Active1 | ASSET | 368.63 | Active | **Inactive** | **No** |
| ASSET_Active2 | ASSET | 207.13 | Active | Active | Yes |
| ASSET_Active3 | ASSET | 219.46 | Active | Active | Yes |
| ASSET_Inactive1 | ASSET | 500.07 | Inactive | Inactive | Yes |
| ASSET_Inactive2 | ASSET | 471.44 | Inactive | Inactive | Yes |
| ASSET_Inactive3 | ASSET | 578.50 | Inactive | Inactive | Yes |
| ***IL-10 (n = 6) RCI threshold = 168.92, the midpoint of the largest gap (sparsest interval) in the dataset's full sorted RCI distribution. largest gap (41.47) falls between IL_10_Active2 (148.19) and IL_10_Inactive3 (189.66); two lower-RCI Inactive sequences (93.61, 133.90) fall below the gap despite their Inactive label.*** | | | | | |
| IL_10_Active1 | IL-10 | 74.92 | Active | Active | Yes |
| IL_10_Active2 | IL-10 | 148.19 | Active | Active | Yes |

| Aptamer ID (Label) | Data Set | RCI | Ground Truth | Predicted | Correct |
|---|---|---|---|---|---|
| IL_10_Active3 | IL-10 | 89.20 | Active | Active | Yes |
| IL_10_Inactive1 | IL-10 | 93.61 | Inactive | **Active** | **No** |
| IL_10_Inactive2 | IL-10 | 133.90 | Inactive | **Active** | **No** |
| IL_10_Inactive3 | IL-10 | 189.66 | Inactive | Inactive | Yes |

**Table S3.** ASSET (under review), IL-10, VEGF, Chloramphenicol, Dengue and Thrombin aptamer set sequences used for 3D Free Energy Landscape Visualizations and Round to Round Enrichment behavior analysis

| NAME | SEQUENCES |
|---|---|
| IL_10 Active_1 | CCCCCGCATCACGCCGTGGTGCGATTGACACAATTGCAAT |
| IL_10 Active_2 | CTCAGCGCGTAGTCGTGTGCGAAACTGCCTTTCCATGGTC |
| IL_10 Active_3 | CGTGACTCGACTCAGGTTTTGCACGGCCTCAGGGTAGCAC |
| IL_10 Inactive_1 | TATAGAGAACTTCTCTCAGTCGAAGCCAAAGAGCATTAAT |
| IL_10 Inactive_2 | ACGATCACTTCTCTCAGTCGGACTAATATTACGGTTAGAA |
| IL-10 Inactive_3 | TGCAATGAGGACTTCTCTCAGTCTAACACAATGTTGTTTA |
| Hoinka 0004 | TAACACTCGATTCTCCTAGCCCGCTAGAAATTCCCCTCCC |
| Hoinka 0008 | TGAGAACTTCTCTCAGTCGGTGGGAGAGTACATCCTAACA |
| Hoinka 00011 | CCCCTTCCAGCGATTACGATCATTGACTCTCAGTCCTGTG |
| Hoinka 00015 | TAGACGAGCACTTCTCTCAGTCGCATTCATTATTTAAATT |
| Hoinka 00029 | TGCAATGAGGACTTCTCTCAGTCTAACACAATGTTGTTTA |

| | |
|---|---|
| Hoinka 00035 | TATAGAGAACTTCTCTCAGTCGAAGCCAAAGAGCATTAAT |
| Hoinka 00057 | TGCCCTGCTTCCCCAGTCTTGTCTCACAAGACTAATTTTA |
| Hoinka 00119 | ACACGACGAGTACAGCTCTCAGTCGAGCGTATTAGCGGAT |
| VEGF SL2-B_Active_1 | CAATTGGGCCCGTCCGTATGGTGGGT |
| VEGF SL12_Active_2 | ATACCAGTCTATTCAATTGGGCCCGTCCGTATGGTGGG |
| VEGF SL1_Inactive_1 | CCAGTCTATTCAATTGG |
| VEGF VEa5_Inactive_2 | ATACCAGTCTATTCAATTGGGCCCGTCCGTATGGTGGGTGTGCTGGCCAGATAGTATGTGCAATCA |
| Chloramphenicol 70Cm6_Active_1 | AUGAAAAGGGCUGGCGAGACAUAUCCGCUGGGCAAUCAGAUUCGGAGCCGCACCACCCUCGAAGUAGACA |
| Chloramphenicol 70Cm9_Active_2 | AGAGCUUGACGGUCCCGAGAGUCGAGCCCAAGCUGACACUGGACCUUUGCGGACCACGUGUUGAUCGUCG |
| Chloramphenicol 70Cm53_Active_3 | GGCACCAAAGCUGAAGUAGCGGGAUAACUCAAAUUACUUUAGGUGUAUGAAGGUGAAACUAGCAAUGAA |
| Chlorampheni | CCCGGCUAGCCGAUACAUCCAUUCGGAACUGCUGACCGUUAGGUGAAUAUCGCCA |

| | |
|---|---|
| col 80Cm51_Inactive_1 | GUCCUACACUGGGAGCUCAUAAGCC |
| Chloramphenicol 80Cm38_Inactive_2 | GCCAAAAGGCAUAAACCACGACGGAGAUUCGGUGGUACGCUGAAUACGUUAGUUAACAACUCCGCUUACGGCAACAGUC |
| Chloramphenicol 80Cm54_Inactive_3 | CUCGGGAUACGGCCUCCCGCGAUCGGACAUUCUCCGGCGGCUCUCAGUACUGGAUAAAAGGGCGGUUUCCCGGCUCAGAG |
| Dengue DENV-3_Active1 | GGAGCUCAGCCUUCACUGCUUUGAUCUCGUGGGGGUGUGUCGCGGGAGACACCAUGGAAUAUAUGGCUGAUUUCAUGUGGGCACCACGGUCGGAUCCAC |
| Dengue DENV-6_Active2 | GGAGCUCAGCCUUCACUGCCGUAGUCGUAUCUCCAUUACCCAGAAUGGUGAUGCGCCGUGAAGAGCGGUUUAGGGAAUUGGCACCACGGUCGGAUCCAC |
| Dengue DENV-2_Inactive1 | GGAGCUCAGCCUUCACUGCUGUCAGAUAACAUGCAUGAUAGACUGAUGAUCGUCCAUGUUUGAAACUGAUCAGUAUCGAGGCACCACGGUCGGAUCCAC |
| Dengue DENV-7_Inactive2 | GGAGCUCAGCCUUCACUGCUUGCUUUUGGGUGCCGUACGCAUUGCGGCAGGGGGAAGAGGAGGGUAGCGACCAGUCGAAGGCACCACGGUCGGAUCCAC |
| Thrombin_16.24_Active_1 | GGGAGAUGCCUGUCGAGCAUGCUGCAUCCGGAUCGAAGUUAGUAGGCGGAGUGGUAGCUAAACAGCUUUGUCGACGGG |
| Thrombin_27.33_Active_2 | GGGAGAUGCCUGUCGAGCAUGCUGGUGCGGCUUUGGGCGCCGUGCUUGACGUAGCUAAACAGCUUUGUCGACGGG |

| | |
|---|---|
| Thrombin_2_Inactive_1 | GGGAGAUGCCUGUCGAGCAUGCUGUACUGGAUCGAAGGUAGUAGGCAGUCACGUAGCUAAACAGCUUUGUCGACGGG |
| Thrombin_5_Inactive_2 | GGGAGAUGCCUGUCGAGCAUGCUGAUAUCACGGAUCGAAGGAAGUAGGCGUGGUAGCUAAACAGCUUUGUCGACGGG |

**Computational Implementation**

All analyses were performed using Python (v3.10)–based workflows that integrated several specialized libraries. Folding and partition function computations were carried out with ViennaRNA (v2.6), while numerical analyses relied on NumPy (v1.26) and SciPy (v1.11). Graph construction and connected component analysis were implemented using NetworkX (v3.2), and multidimensional scaling (MDS) along with other dimensionality reduction techniques utilized Scikit-learn (v1.3). Visualizations were generated using Plotly (v5.18) and Matplotlib (v3.8). Thermodynamic calculations followed the ViennaRNA partition function formalism[45], modeling solution-phase free energies under standard conditions. All derived measures were computed directly from sequence-derived ensembles, ensuring that any observed classification signals reflected intrinsic folding organization rather than being influenced by external training labels.

**Supplemental References**


1. Barros M, Kasirajan G, Jones A, Schlichting A, Ruiz-Ciancio D, Lin LH, Narayan C, Veeramani S,Thiel KW, Kennedy GC, Darcy I, Thiel W. (2025). ASSET: A Framework for Decoding Aptamer Specificity of an Enriched Library by Next-Generation Sequencing of Experimental Samples. bioRxiv [Preprint]. Aug 30:.08.27.672406. doi: 10.1101/2025.08.27.672406. PMID: 40909745; PMCID:PMC12407934.
2. Levay A, Brenneman R, Hoinka J, Sant D, Cardone M, Trinchieri G, Przytycka TM, Berezhnoy A. (2015). Identifying high-affinity aptamer ligands with defined cross-reactivity using high-throughput guided systematic evolution of ligands by exponential

enrichment. Nucleic Acids Res. Jul 13;43(12): e82. doi: 10.1093/nar/gkv534. Epub May 24. PMID: 26007661; PMCID: PMC4499151.

3. Kaur H, Yung LY. (2012). Probing high affinity sequences of DNA aptamer against VEGF165. PLoS One.;7(2):e31196. doi: 10.1371/journal.pone.0031196. Epub 2012 Feb 16. PMID: 22359573; PMCID: PMC3281051.

4. Burke DH, Hoffman DC, Brown A, Hansen M, Pardi A, Gold L (1997). RNA aptamers to the peptidyl transferase inhibitor chloramphenicol. Chem Biol. 1 Nov;4(11):833-43. doi: 10.1016/s1074-5521(97)90116-2. PMID: 9384530.

5. Thevendran R, Rogini S, Leighton G, Mutombwera A, Shigdar S, Tang TH, Citartan M. (2023). The Diagnostic Potential of RNA Aptamers against the NS1 Protein of Dengue Virus Serotype 2. Biology (Basel). May 15;12(5):722. doi: 10.3390/biology12050722. PMID: 37237536; PMCID: PMC10215423.

6. Kubik MF, Stephens AW, Schneider D, Marlar RA, Tasset D. (1994). High-affinity RNA ligands to human alpha-thrombin. Nucleic Acids Res.Jul 11;22(13):2619-26. doi: 10.1093/nar/22.13.2619. PMID: 7518917; PMCID: PMC308218
